# LINE-1 retrotransposons regulate the exit of human pluripotency and early brain development

**DOI:** 10.1101/2025.01.17.633315

**Authors:** Anita Adami, Raquel Garza, Patricia Gerdes, Pia A. Johansson, Fereshteh Dorazehi, Symela Koutounidou, Laura Castilla-Vallmanya, Diahann A.M. Atacho, Yogita Sharma, Jenny G. Johansson, Oliver Tam, Agnete Kirkeby, Roger A. Barker, Molly Gale-Hammell, Christopher H. Douse, Johan Jakobsson

## Abstract

Long interspersed nuclear element 1 (L1) retrotransposons represent a vast source of divergent genetic information. However, mechanistic analysis of whether and how L1s contribute to human developmental programs is lacking, in part due to the challenges associated with specific profiling and manipulation of human L1 expression. Here we show that thousands of hominoid-specific L1 integrants are expressed in human induced pluripotent stem cells and cerebral organoids. The activity of individual L1 promoters is surprisingly divergent and correlates with an active epigenetic state. Efficient on-target CRISPRi silencing of L1s revealed nearly a hundred co-opted L1-derived chimeric transcripts and L1 silencing resulted in changes in neural differentiation programs and reduced cerebral organoid size. Together, these data implicate L1s and L1-derived transcripts in hominoid-specific CNS developmental processes.

## Introduction

The human genome consists of at least 50% transposable elements (TEs) that have been incorporated and expanded by waves of retrotransposition, resulting in significant interspecies and interindividual variation in genomic composition^1,2,3,4^. Due to their ability to mobilize, TEs pose a potential threat to genomic integrity^5^. TEs are therefore assumed to be transcriptionally silenced in mammalian cells via epigenetic mechanisms such as DNA methylation^6,7,8,9,10^. However, recent studies have begun to challenge this simplistic view, by demonstrating that some TEs are selectively expressed in various mammalian cell types during early development where they may contribute to important cellular functions^11,12,13,14,15^.

Long interspersed nuclear elements–1 (L1s) are the most abundant and only autonomously mobilizing family of TEs in humans, accounting for ∼17% of human genomic DNA^4,16,1^. Because L1s colonized the human genome in different waves via a copy-and-paste mechanism, it is possible to approximate the evolutionary age of each individual L1 element and assign them to chronologically-ordered subfamilies^17^. Hundreds of thousands of L1s are primate-specific and thousands are human-specific^18^. L1s are transcribed from a CpG-rich internal 5′ RNA polymerase II promoter as a bicistronic mRNA encoding two proteins, ORF1p and ORF2p, which are essential for L1 mobilization^19,20,21–23^. Notably, the L1 promoter is bidirectional, and in evolutionarily young L1s, the antisense transcript encodes a small peptide, ORF0, whose function is poorly characterized^24^. L1 antisense promoters can also give rise to chimeric transcripts and act as alternative promoters for non-canonical isoforms of protein-coding genes^8,25,26^. L1s are highly expressed during early mouse development, where they regulate global chromatin accessibility and influence developmental potency and self-renewal^11,12^. In particular, L1s are expressed in the developing brain, where they regulate the rate of neuronal differentiation and maturation^27–29^. The mechanism by which L1s exert these roles remains debated but appears to be independent of retrotransposition^11,12,27,28^. Rather, L1s may influence transcriptomes via *cis*-acting mechanisms, acting as enhancers or alternative promoters, or via *trans*-acting mechanisms, where they act as regulatory non-coding RNAs^30^. However, the functional relevance of L1 expression in early human development remains poorly understood. The intact L1 loci expressed during mouse development are not present in the human genome, where other primate and human-specific L1 loci are found^4^. Human L1s have integrated at different sites in the genome, and their expression is at least to some extent controlled by different regulatory elements^31^. Thus, L1 expression in human cells is likely to be fundamentally different from that in mouse and may also be functionally very distinct.

In this study, we used a combination of short and long-read RNA-seq approaches coupled with targeted cleavage and nuclease release (CUT&RUN) epigenomic profiling, long-read DNA methylation analysis, and tailored bioinformatic approaches to demonstrate that L1-derived transcripts are highly expressed in human induced pluripotent stem cells (hiPSCs) and human cerebral organoids and that their expression correlates with the presence of active histone marks and the absence of DNA methylation. We applied an optimized CRISPR interference (CRISPRi) based system that allows efficient and specific silencing of L1 expression in hiPSCs and organoids and found that L1s exert profound *cis*-acting effects on gene expression by acting as alternative promoters for almost 100 protein-coding genes and long non-coding RNAs (lncRNAs). This L1-mediated *cis*-activity is not essential for the maintenance of pluripotency in human iPSCs but is important for the regulation of neural differentiation in cerebral organoids. L1 silencing reduces the size of cerebral organoids at timepoints corresponding to the proliferation of neural progenitors. In summary, these results demonstrate that L1s are wired into gene regulatory networks in human pluripotent stem cells and provide a layer of primate- and human-specific transcriptome complexity that influences neural differentiation and may have contributed to the evolution of the human brain.

## Results

### Evolutionarily young L1s are highly expressed in hiPSCs

The human genome contains approximately half a million individual L1 copies, most of which are transcriptionally inactivated due to 5’ truncations and the accumulation of mutations and deletions^4^. Only L1s with intact 5’ promoter regions are capable of driving L1 transcription, and our genome contains approximately 6,000 full-length (>6-kbp) L1 elements (Figure 1a). We characterized the expression of individual full-length L1 elements in two hiPSC lines (hereafter referred to as hiPSC1 and hiPSC2, see Methods for details) (Supplementary Figure 1a). We generated bulk RNA sequencing data using an in-house 2 x 150 base pair (bp), polyadenylate (poly-A) enriched stranded library preparation with a reduced fragmentation step to optimize library insert size for L1 analysis (Figure 1a). We obtained an average of 58 million reads per sample. Due to their repetitive nature, a large proportion of reads originating from L1s are expected to ambiguously map to the reference genome^26,32^. To avoid an inflated signal due to multimapping reads, we used a unique mapping approach to investigate the expression of individual L1 elements (Figure 1a). This bioinformatic approach yielded an average of 85.9% uniquely mapped reads per sample. By using these uniquely mapped reads it is possible to investigate the transcriptional status of most individual L1 loci, except for some of the most evolutionarily recent integrants, as well as polymorphic L1 alleles that are not present in the hg38 reference genome^26^.

**Figure 1.**
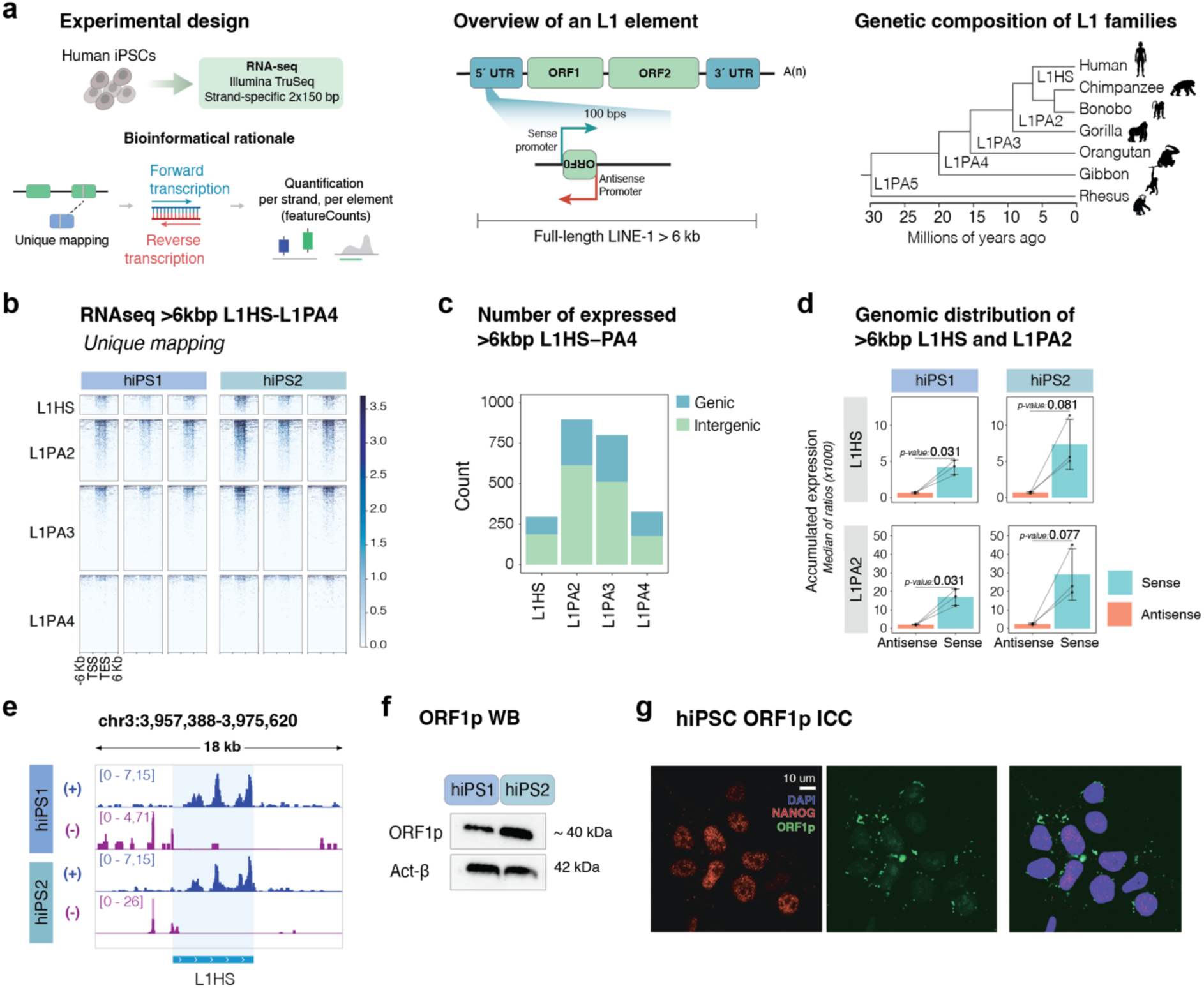
L1s are highly expressed in human iPSCs. a) Left: Schematic of the experimental design (top panel) to profile L1 expression in human induced pluripotent stem cells (hiPSCs). Bioinformatic rationale (lower panel) to quantify L1 expression by using a unique mapping approach to quantify stranded expression of unique L1 loci. Middle: Overview of a full-length L1 element with an intact 5’UTR and two long open reading frames (ORF1 and ORF2). The zoom in shows the bidirectional promoter encoding the L1 body in sense and ORF0 in antisense. Arrows indicate the sense and antisense promoters. Right: Phylogenetic tree showing the evolutionary age of different L1 subfamilies. b) Expression [reads per kilobase per million mapped reads (RPKM)] of full-length (>6kbp), evolutionarily young L1s in two hiPSC lines. n= 3 technical replicates. c) Number of L1s belonging to primate-specific L1 subfamilies (L1HS to L1PA4) expressed in hiPSCs. d) Normalised read counts in in antisense (red) and sense (blue) and of full-length L1HS (top) and L1PA2 (bottom) per sample (n=3 technical replicates, t-test). Bar representing mean normalized expression, error bars showing mean +/- standard deviation. e) Genome browser tracks showing normalized expression (RPKM) of an L1HS element in two hiPSC lines (dark blue: forward transcription; purple: reverse transcription). f) Western blot (WB) of ORF1p in two hiPSC lines (top), and actin-β (bottom) as loading control protein. g) Immunocytochemistry of the pluripotency marker NANOG (red) and the L1-derived protein ORF1p (green) in hiPSCs. DAPI nuclear staining (blue) is included in the overlay.

We found that many individual full-length L1 loci were highly expressed in hiPSCs. These L1s belonged predominantly to primate-specific families (Figure 1b-e), including both hominoid-specific elements (L1PA2 to L1PA3) and human-specific elements (L1HS). We detected the expression of 2,323 unique loci of evolutionarily young L1s (Figure 1b-c). The RNA-seq signal was present throughout the L1 body, with a slight enrichment at the 3′ end, suggesting that transcription of most L1s terminates in the internal L1 polyadenylation signal (Figure 1b&e). Most of the transcription was in the same orientation as the L1s (in sense, Figure 1d, Supplementary Figure 1b). This suggests that most L1 transcripts originate from the L1 promoter and are not a consequence of readthrough or bystander transcription. We also found evidence for activity of the antisense L1 promoter, resulting in transcription extending into the upstream flanking genome (Figure 1e, Supplementary Fig 1b).

L1s encode open reading frames that lead to the translation of peptides, including the RNA binding protein ORF1p. Western blot (WB) analysis confirmed that ORF1p is highly expressed in hiPSCs (Figure 1f). Immunocytochemistry (ICC) of ORF1p in hiPSCs showed localization to cytoplasmic, perinuclear puncta, consistent with what has been previously described in other human cell lines (Figure 1g)^33^. Taken together, these results demonstrate that L1 elements are highly expressed in human pluripotent stem cells.

### Transcription of unique L1 loci correlates with their epigenetic status

The RNA and protein analyses show that L1s are highly expressed in hiPSCs, raising the possibility that the epigenetic status of L1 promoters is associated with an active state in this cell type. To address this question, we performed cleavage under targets and release using nuclease (CUT&RUN) epigenomic analysis^34^ to determine whether the histone mark H3K4me3, which is associated with active promoters, is present on evolutionarily young L1s in iPSCs (Figure 2a, Supplementary Figure 2a). An additional advantage of H3K4me3 profiling is that the signal of these histone modifications spreads to the unique flanking genomic context, allowing accurate identification of individual transcriptionally active L1 promoters^26^. The resulting sequencing data was uniquely mapped, followed by peak calling and intersection with full-length L1s. The CUT&RUN analysis identified 133 high-confidence H3K4me3 peaks located at the 5’end of evolutionarily young full-length L1s, and most of these loci were confirmed to be expressed in the bulk RNA-seq dataset (Figure 2b-c).

**Figure 2.**
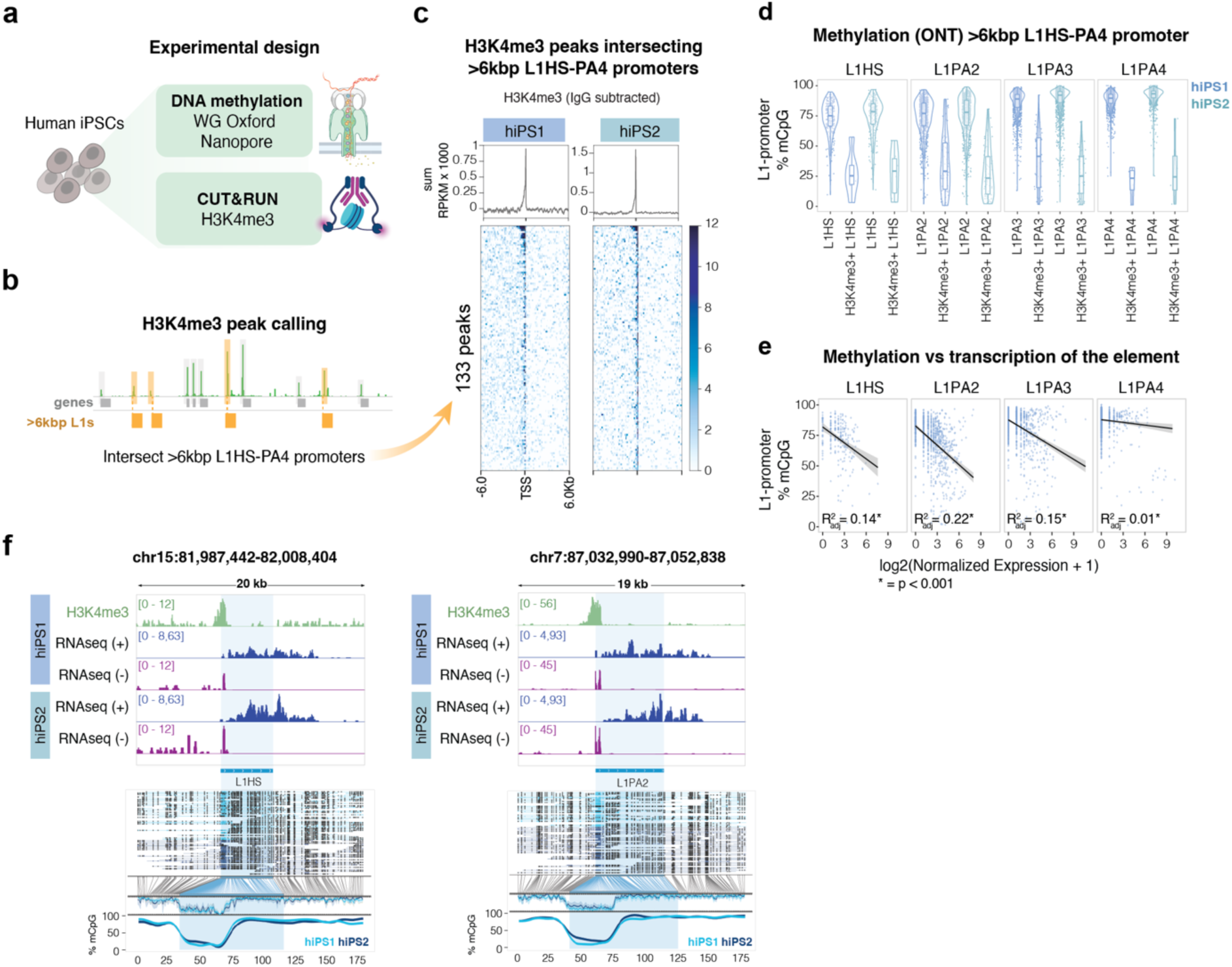
The epigenetic profile over L1 loci correlates with their expression. a-b) Schematic of the CUT&RUN and DNA methylation analysis workflow. c) H3K4me3 peaks over the promoter of young (L1HS-PA4) >6kbp L1s (n=2 biological replicates) (RPKM normalized). TSS = transcription start site. Profile plot at the top showing the summed signal. d) Violin plots showing DNA methylation levels (ONT DNA sequencing data) over the promoter of full-length (>6kbp), evolutionary young (L1HS-L1PA4) L1 elements in two hiPSC lines, including H3K4me3 peak called L1s. e) Scatter plots showing a negative correlation between normalized expression of individual FL-L1s (x-axis; normalized counts by gene sizeFactors as calculated by DESeq2 (i.e. median of ratios)) and the percentage of methylated CpG sites at their promoters (y-axis) of full length (>6kbp). R2 and p-value of fitted linear models between methylation and expressed per L1 subfamily. f) Genome browser tracks showing from top to bottom: H3K4me3 mark at the L1 promoter (RPKM normalized), transcription of the element (dark blue: forward transcription, purple: reverse transcription. RPKM normalized). ONT DNA reads in the region, black dots indicating methylated CpG sites, and methylation coverage of the L1 elements at the bottom (light blue hiPSC1, dark blue: hiPSC2).

Transcriptional silencing of L1s has previously been associated with the presence of CpG DNA methylation at the L1 promoter^8,9^. To investigate whether L1 expression in hiPSCs correlates with the absence of DNA methylation, we performed genome-wide methylation profiling using long-read sequencing from Oxford Nanopore Technologies (ONT) (Figure 2a). This analysis revealed that the DNA methylation level of the L1 promoter negatively correlated with their evolutionary age. Promoters of human-specific L1 elements were hypomethylated when compared to more ancient subfamilies (Figure 2d). Notably, L1s that carried a H3K4me3 mark displayed very low levels of DNA methylation over the L1 promoter and the lack of DNA methylation at individual L1 loci correlated with the RNA expression at the same loci (Figure 2d-e). For example, we found several instances where the L1 promoter was hypomethylated in hiPSCs and where this correlated with the presence of H3K4me3 and RNA-seq signal (Figure 2f). These results demonstrate that the epigenetic status of each individual L1 promoter in hiPSCs, including the absence of DNA methylation and the presence of H3K4me3, correlates with the transcriptional activity of individual loci.

### Efficient, on-target L1 silencing using CRISPRi

To investigate the functional role of L1s in human pluripotency and early human brain development, we optimized a lentiviral CRISPR interference (CRISPRi) system^35^ to transcriptionally silence L1s in hiPSCs (Figure 3a-b). To achieve specific and efficient transcriptional silencing, we tested several gRNA designs (see Methods for details). Ultimately, we selected two distinct 20 bp gRNAs that target the 5’UTR of young, full-length L1s close to their transcription start site (TSS)^36^ which, when co-expressed with a Krüppel-associated box transcriptional repressor domain fused to a catalytically inactive Cas9 (KRAB-dCas9), efficiently silenced L1 transcription (Figure 3a-e, Supplementary Figure 3a). As a control, we transduced hiPSCs with the same lentiviral CRISPRi system but using a non-targeting control gRNA (see Methods for details).

**Figure 3.**
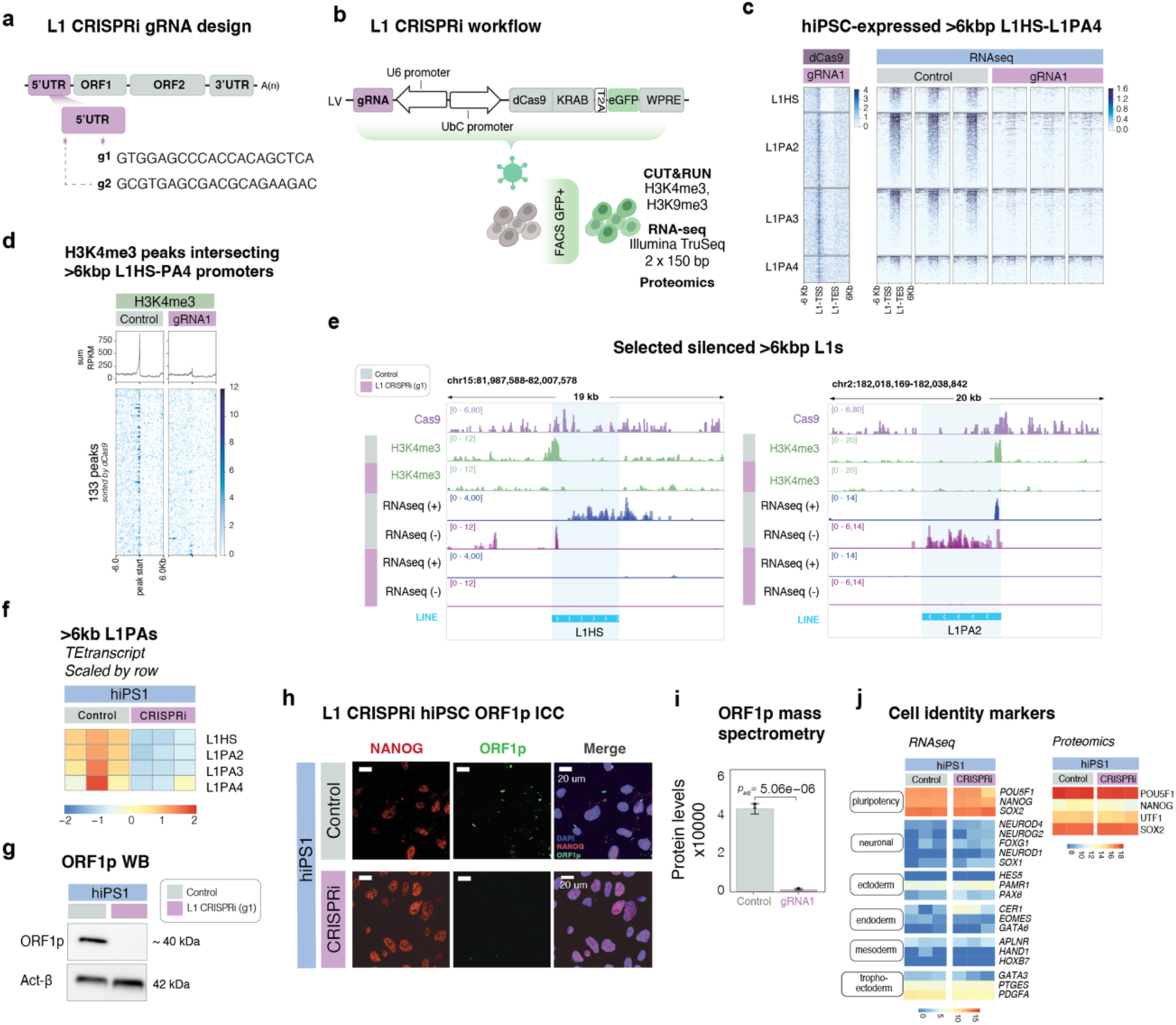
CRISPRi-based silencing of L1s in hiPSCs does not affect human pluripotency. a) Schematic of the gRNA target sites within the full-length L1. The gRNAs were designed to target the 5’ UTR of evolutionarily young, >6kbp L1s. b) Schematic of the CRISPRi workflow and downstream analyses. c) CUT&RUN analysis of dCas9 (gRNA1 – Control signal) (left) and bulk RNA seq data showing the normalized expression of uniquely mapped, full-length (>6kbp), evolutionary young (L1HS to L1PA4) L1s in L1-CRISPRi vs. control hiPSCs (all tracks RPKM normalized). d) H3K4me3 peaks over the promoter of young (L1HS-PA4) >6kbp L1s in L1-CRISPRi vs. control hiPSCs. Profile plot at the top showing the summed signal. e) Genome browser tracks illustrating dCas9 enrichment (gRNA1 – Control signal) and H3K4me3 loss over L1 elements’ promoter, and the loss of expression in L1-CRISPRi hiPSCs vs. control (dark blue: forward transcription; purple: reverse transcription) (All tracks RPKM normalized). f) Expression of L1 families analysed using tEtranscripts in L1-CRISPRi vs. control hiPSCs (heatmap showing normalized expression, scaled by row). g) Western blot (WB) of ORF1p (top) and actin-β (bottom) in L1-CRISPRi hiPSCs vs. control. h) Immunostaining of the pluripotency marker NANOG (red) and the L1-derived protein ORF1p (green) in L1-CRISPRi vs. control hiPSCs. DAPI nuclear staining in blue. i) Mass spectrometry (MS) data showing changes (p_adj_) in ORF1p levels in L1-CRISPRi hiPSCs. See Methods for statistical analysis details. j) Left panel: Heatmap showing log2 normalized expression (RNA-seq) of pluripotency and differentiation markers in L1-CRISPRi vs. control hiPSCs. Right panel: Heatmap of MS data showing expression of pluripotency markers in L1-CRISPRi vs. control hiPSCs (heatmap showing log2 normalized expression).

To evaluate the efficiency and on-target specificity of the lentiviral CRISPRi system, we performed an in-depth epigenetic profiling of the L1-CRISPRi hiPSCs and control transduced cells. We performed CUT&RUN analysis of the dCas9 protein, which is expected to be recruited to the gRNA target site, as well as the active histone mark H3K4me3 and the repressive histone mark H3K9me3 (Figure 3d; Supplementary Figure 3b). Using unique mapping, we observed clear enrichment of dCas9 protein at the 5’ UTR of evolutionarily young, full-length (>6kbp) L1s in L1-CRISPRi hiPSCs (Figure 3c). This was accompanied by a loss of H3K4me3 and gain of H3K9me3 at gRNA binding sites (Figure 3d, Supplementary Figure 3b), confirming efficient on-target engagement of the L1-CRISPRi system at the promoter of evolutionarily young L1s. To further characterize the CRISPRi system, we examined other (non-L1) genomic loci where a dCas9 peak was detected (n=272 CUT&RUN peaks) (Supplementary Figure 3e-f). Importantly, these peaks were also detected in cells expressing the non-targeting gRNA, and the levels of H3K4me3, H3K9me3 and transcription were unchanged between L1-CRISPRi and control cells at these sites, strongly implying that these are technical artefacts of the dCas9 CUT&RUN experiment, which required a crosslinking approach, rather than off-target effects associated with the L1 gRNAs (Supplementary Figure 3f).

To investigate the transcriptional consequences of L1-CRISPRi transduction we performed RNA-seq and analyzed the expression of unique L1 loci (Figure 3c&e, Supplementary Figure 3c). We found that the expression of evolutionarily young full-length L1 elements was almost completely silenced in L1-CRISPRi hiPSCs, including both in-sense transcription and antisense promoter activity (Figure 3c&e, Supplementary Fig 3c-d). To quantify the extent of L1 silencing, we used the TE-oriented read quantification software TEtranscripts^37^. This approach confirmed a global transcriptional silencing of evolutionarily young L1 families (Figure 3f, Supplementary Figure 3g), leading to complete loss of the L1-derived protein ORF1p in the L1-CRISPRi hiPSCs assessed with WB, immunocytochemistry and quantitative mass spectrometry (MS) (Figure 3g-i, Supplementary Figure 3h-i). We confirmed efficient L1 silencing using the lentiviral CRISPRi system at both the RNA and protein levels using both of our two gRNAs in the two different hiPSC lines (Supplementary Figure 3c-d&3g-i). Taken together, these data demonstrate efficient and specific on-target activity of the L1-CRISPRi system.

### L1 expression is not essential for the maintenance of human pluripotency

Two previous studies have suggested that L1s may play a critical role in early mouse development and that L1 expression is essential for maintaining pluripotency^11,12^. Recently, similar observations have also been made in human pluripotent stem cells^38^. However, we found that L1-CRISPRi hiPSCs remained proliferative and exhibited a morphology characteristic of human pluripotent stem cells (Figure 3h, Supplementary Figure 3i-j). Consistent with this, RNA-seq analysis confirmed that L1-CRISPRi hiPSCs maintained high expression of pluripotency-related genes, such as *POU5F1*, *NANOG* and *SOX2* while genes associated with differentiation of all other lineages (ecto-, endo-, meso-, trophoecto-dermal markers) remained transcriptionally silent (Figure 3j, Supplementary Figure 3k). We also verified the expression of pluripotency-related proteins in L1-CRISPRi using the mass spectrometry data and found high levels of expression of *POU5F1, NANOG, UTF1* and *SOX2* (Figure 3j, Supplementary Figure 3k). The expression of NANOG was further verified by immunostaining (Figure 3h, Supplementary Figure 3i). These data show that the L1-CRISPRi hiPSCs remain in a proliferative pluripotent state, demonstrating that L1 transcription is not essential for the maintenance of human pluripotent stem cells.

### L1s influence the expression of protein coding genes and long non-coding RNAs *in cis*

To investigate the transcriptional consequences of L1 silencing in hiPSCs, we performed differential gene expression analysis comparing L1-CRISPRi transduced cells to the control cells (Figure 4a). We found that 99 genes were significantly downregulated (p-adj <0.05, log2, fold-change >1) in L1-CRISPRi hiPSCs, while only 6 genes were upregulated. Most of the downregulated genes (61/99) were in the vicinity (≤50kbp) of a CRISPRi-silenced L1 (Figure 4b), showing that the transcriptional silencing of L1s impacts mainly on nearby gene expression. Overall, this indicates that most of the gene expression changes in the L1-CRISPRi hiPSCs are direct *cis*-related effects of the L1 transcriptional silencing, while any downstream effects appear to be minor. Among the downregulated genes, 67 were non-coding RNAs, mostly lncRNAs, while 32 were protein-coding genes (Figure 4b&c). This transcriptional response in L1-CRISPRi hiPSCs was highly reproducible with the second L1-targeting gRNA as well as in the second hiPSC line, suggesting that these 99 transcripts are likely to be directly regulated by L1s *in cis* in hiPSCs (Figure 4c and Supplementary Figure 4b).

**Figure 4.**
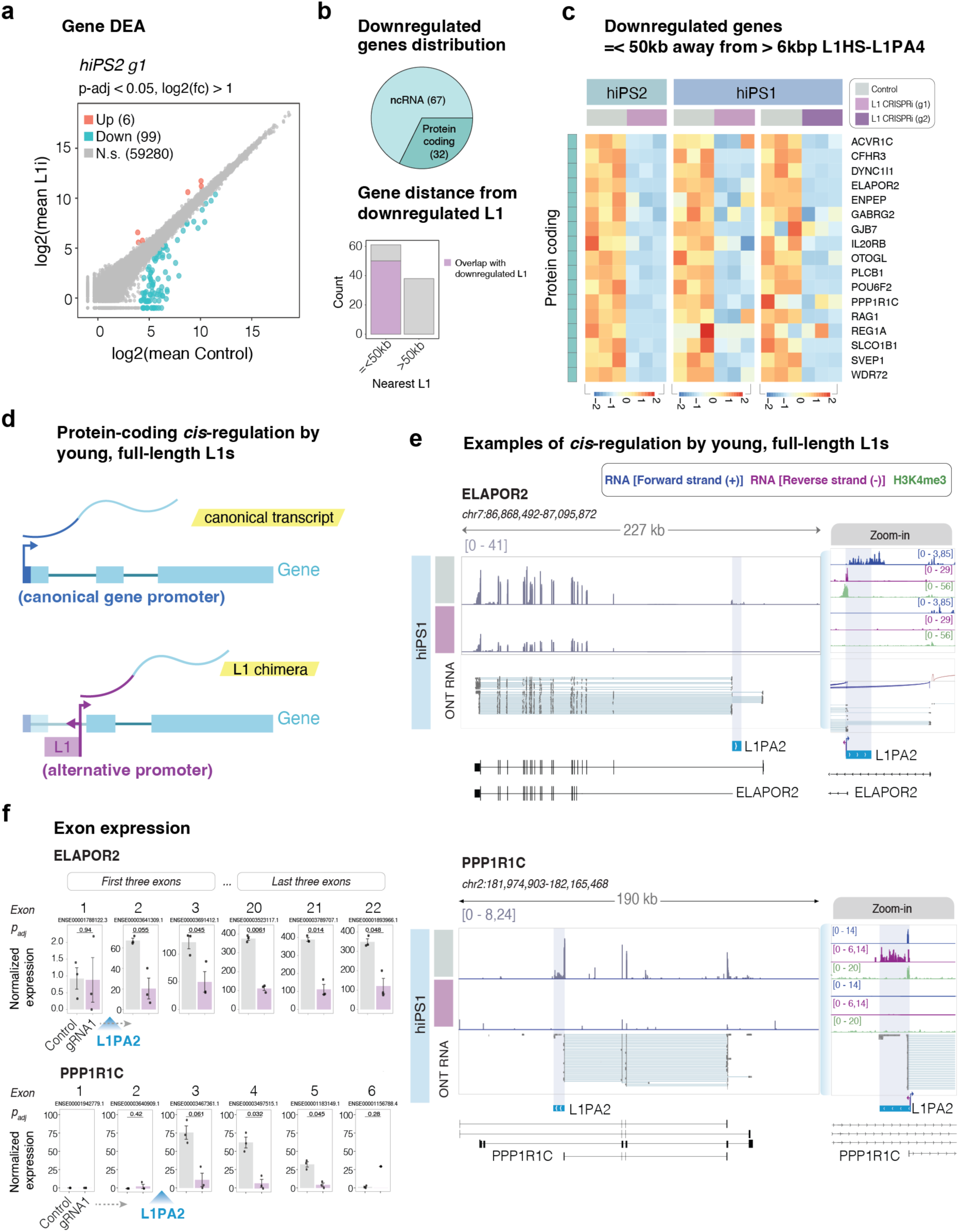
L1s drive the expression of protein coding genes and long non-coding RNAs in hiPSCs. a) Scatterplot showing mean gene expression in L1-CRISPRi (y-axis) and control (x-axis) hiPSCs and summary of the differential expression analysis (DEA) (Bulk RNA-seq, n = 3 replicates / condition). Blue dots = significantly downregulated genes; red dots = significantly upregulated genes; grey dots = non-significant (DESeq2: Wald test; p-adj <0.05; log2(foldChange) >1). b) Top panel: Distribution between protein-coding and non-coding downregulated genes. Lower panel: number of downregulated genes split by their distance from the nearest FL-L1. In purple: number of genes that overlap with a FL-L1 (intragenic). c) Heatmap showing all the normalized expression of downregulated protein-coding genes that overlap with a FL-L1 (n= 3 replicates / condition, heatmap showing log2 normalized expression). d) Schematic showing the expression of a canonical transcriptional isoform from the canonical gene promoter vs. expression of an L1-driven alternative transcriptional isoform. e) Genome browser tracks showing the expression of L1-driven alternative transcriptional isoforms at the *ELAPOR2* and *PPP1R1C* loci in human iPSCs, and its silencing upon L1 CRISPRi. Left panel showing normalized transcription (RPKM) in L1-CRISPRi and control hiPSCs ond ONT long-read direct RNA from control hiPSCs showing the different isoforms. Right panel with a zoom-in to the L1 elements showing from top to bottom: bulk RNA-seq data (dark blue: forward transcription, purple: reverse transcription), H3K4me3 CUT&RUN (green, control vs L1-CRISPRi hiPSCs), and ONT direct RNA reads from control hiPSCs. f) Normalized exon expression between L1-CRISPRi and control hiPSCs of the first three and last exons of ELAPOR2, and all the exons of PPP1R1C (p_adj_ DESeq2: Wald test, n = 3 replicates per condition). Intronic L1 position is indicated in blue.

L1s can act as alternative promoters for protein-coding genes using the activity of the L1 antisense promoter^8,25,26^ (Figure 4d). Focusing on the downregulated protein-coding genes, we indeed found several examples of this phenomenon where genes on the opposite strand to a nearby L1 were transcriptionally silenced upon L1-CRISPRi (Figure 4d-e). These observations suggest that the antisense L1 promoter drives expression of an alternative isoform of these protein coding genes. Direct long-read ONT RNA-sequencing analysis (Supplementary Figure 4c) of hiPSCs confirmed the existence of L1-fusion transcripts in these genes (Figure 4e, Supplementary Figure 4d-e). For example, an L1PA2 provides an alternative promoter for a hominoid isoform of *ELAPOR2*, a gene involved in the BMP signaling pathway^39^ and another L1PA2 provides an alternative hominoid promoter for *PPP1R1C*, which is part of a major serine/threonine phosphatase (Figure 4e). The direct long read RNA-seq analysis confirmed that the majority of *ELAPOR2* transcripts start in the L1 element and are then spliced into the conserved downstream exons (Figure 4e). The L1PA2 promoter in *PPP1R1C* is driving almost all detected expression of this gene in hiPSCs (Figure 4e). Analysis of exon usage of these two genes in hiPSCs confirmed that exons downstream of the L1 insertion are highly expressed and downregulated by L1-CRISPRi transduction, while exons upstream of the L1 insertion are lowly expressed and not affected by L1-CRISPRi (Figure 4f), confirming that L1-promoter activity is responsible for most of the transcription of these two genes in hiPSCs. We also found that the majority of the downregulated lncRNAs use an L1 antisense promoter to drive their expression (Supplementary Figure 4b&d). One such example is *LINC00648*, a lncRNA involved in various types of cancer^40,41^. *LINC00648* is highly expressed in hiPSCs and uses an L1PA2 as its promoter. Upon L1-CRISPRi, the expression of this lncRNA is completely shut down (Supplementary Figure 4e) together with the L1PA2 element, confirming the L1 as the driver of transcription.

Taken together, the L1-CRISPRi RNA-seq analysis revealed nearly a hundred genes that depend on L1 promoter activity in human pluripotent cells. All these L1s represent hominoid or human-specific insertions, demonstrating that these elements provide a hominoid-specific layer of transcriptome complexity during early human development. Notably, several of these genes, including *ELAPOR2, PPP1R1C* and *PLCB1* (Figure 4c&e), are genes involved in brain development^39,42,43^, suggesting a potential role for *cis*-acting L1s in hominoid brain speciation. Consistent with this, RNA-seq and CUT&RUN analysis of human fetal forebrain tissue^26^ confirmed the presence of several of these L1-driven transcript isoforms in the developing human brain, suggesting that they may play a functional role *in vivo* (Supplementary Figure 4f).

### Expression of L1 loci in in cerebral organoids is variable and DNA methylation dependent

To study the role of L1s in human brain development, we generated human cerebral organoids, as a model for human brain development in a 3D setting (Figure 5a). We cultured hiPSC-derived unguided cerebral organoids for 15 days. At this time point the organoids are mostly composed of neural rosettes, as determined using brightfield imaging and ZO1/PAX6 immunostaining (Figure 5b, Supplementary Figure 5a), representing an early stage of human brain development when neural progenitor cells (NPCs) are expanding^44,45^. Using snRNA-seq analysis, we confirmed that most of the cells displayed a transcriptome characteristic of NPCs, while a minority of the cells expressed genes associated with early neurons (Figure 5c, Supplementary Figure 5b). We found no evidence for the presence of pluripotent cells in the organoids, confirming that this is a suitable model system to study the exit from the pluripotent stage into a committed neural progenitor cell.

**Figure 5.**
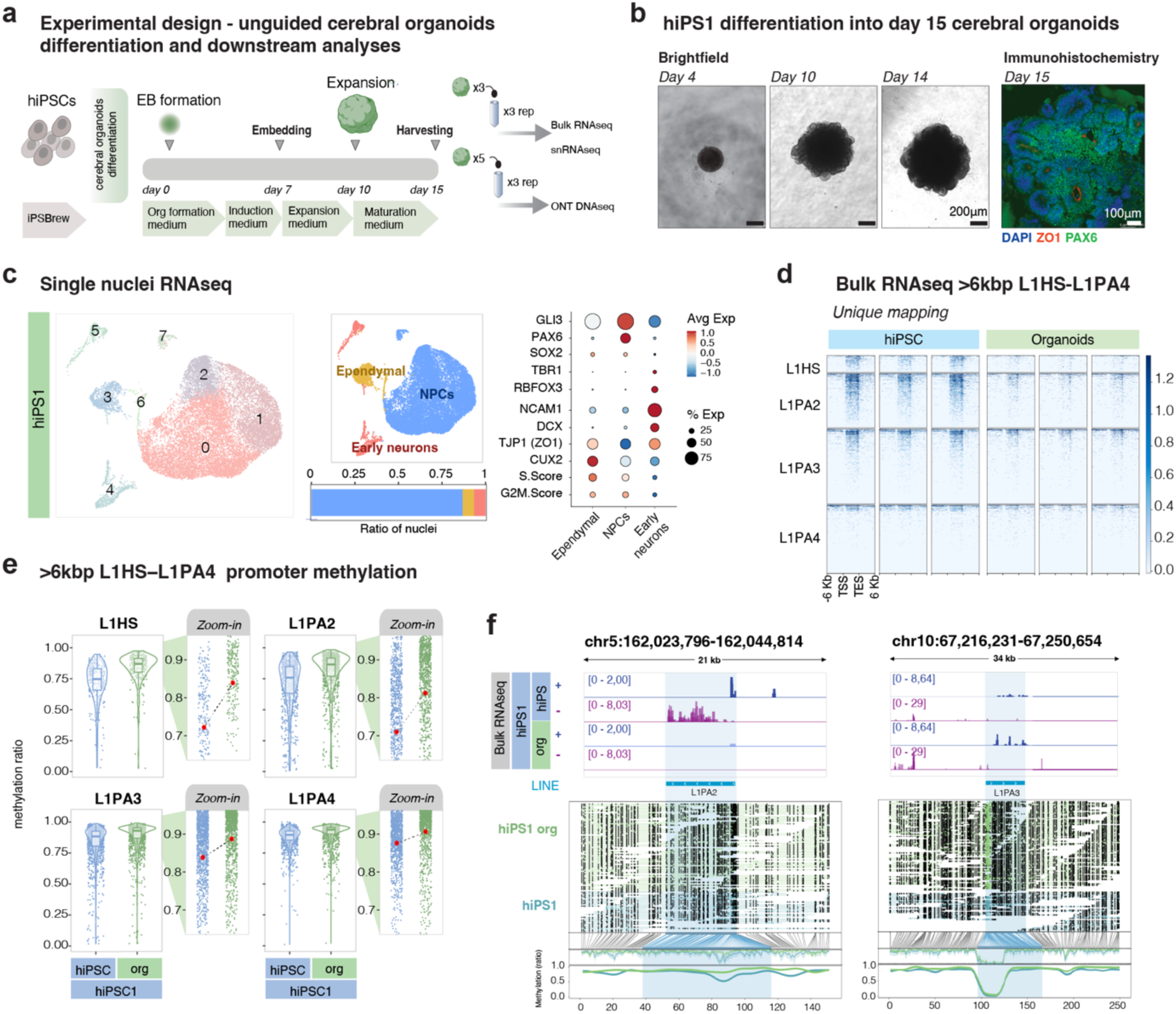
Expression of evolutionary young full-length L1s in cerebral organoids. a) Schematic of the workflow for the generation of unguided cerebral organoids and downstream analyses. b) Brightfield images of differentiating cerebral organoids at different time points and immunohistochemistry on day 15 organoids for ZO1 (red) and PAX6 (green). DAPI in blue. c) Upper: UMAP showing cell types found in day 15 unguided cerebral organoids. Middle: bar plot showing the percentage of the cell type composition of the cerebral organoids at day 15 of differentiation. Bottom: dot plot displaying selected neuronal and NPC markers used to characterize the cell clusters (dot size showing the percentage of cells expressing the gene, color indicates average expression in each cell type). d) Expression of uniquely mapped evolutionary young FL-L1s in hiPSCs and day 15 cerebral organoids. e) Violin plots of the methylation status over the promoter of L1HS-L1PA4 >6kbp L1s in hiPSCs vs. day 15 cerebral organoids. Zoom in panels indicating mean methylation levels (red dot) per condition per subfamily. f) Genome browser tracks showing normalized (RPKM) of L1s in hiPSCs and day 15 cerebral organoids (dark blue: forward transcription, purple; reverse transcription) and ONT DNA reads across day 15 organoids and hiPSCs, black dots indicating methylated CpGs, and methylation coverage of the L1 elements at the bottom (green: organoids, blue: hiPSCs).

To investigate L1 expression, we performed bulk RNA sequencing, which revealed that L1s were expressed in cerebral organoids, albeit at lower levels compared to hiPSCs (Figure 5d, Supplementary Figure 5c). The reduced L1 expression correlated with an overall increase in DNA methylation at L1 promoters in organoids compared to hiPSCs (Figure 5e). This is consistent with the restoration of DNA methylation during differentiation^46^. However, of the 2323 L1s expressed in hiPSCs, 750 L1s were completely silenced in organoids, while the expression of 1573 loci could still be detected (Supplementary Figure 5d). This difference was also reflected in the DNA methylation status, where L1s expressed in organoids showed lower levels of DNA methylation at the L1 promoter than the silenced loci (Supplementary Figure 5e). This suggests that when analyzing the epigenetic status of L1s, it is not possible to treat them as a single uniform family. Rather, each locus seems to be regulated in an independent manner, as previously suggested^9,10,47,48^. We found fully methylated L1 loci that were transcriptionally silent in organoids, as well as fully demethylated loci in organoids that were expressed at similar levels as in hiPSCs (Figure 5f). This is surprising, as the different L1 loci share the same regulatory sequences, but mirrors observations we have made previously in the developing and adult human brain^26^. Taken together, these results demonstrate that at the global level, the expression of L1s depends on the cellular context during early human development and neural differentiation, but that epigenetic state and transcriptional activity of each individual L1 element is surprisingly variable.

### L1s regulate early neural differentiation in cerebral organoids

To investigate whether L1 expression plays a role in the exit from human pluripotency and early neural differentiation, we generated L1-CRISPRi organoids. L1 silencing did not affect the formation of the organoids, which displayed characteristic neural rosettes after 15 days of differentiation, as well as expression of ZO1 and PAX6 (Figure 6a-c, Supplementary Figure 6a-b). To assess the cellular and molecular consequences of L1-CRISPRi inhibition on human cerebral organoids, we used snRNA-seq (Figure 6d-f). High-quality data were generated from a total of 66206 cells, including 39181 from L1-CRISPRi organoids (two gRNAs, two cell lines; in total 11 libraries made out of 5 organoids each) and 27025 from control organoids (LacZ-gRNA, two cell lines in total; 8 libraries made out of 5 organoids each). We performed an unbiased clustering analysis to identify and quantify the different cell types present in the organoids. Eight separate clusters of cells were identified, with the majority of cells being non-proliferating NPCs (cluster 0) and proliferating NPCs (cluster 1), as well as clusters containing newborn neurons at different stages of maturation (Figure 6d-e). All clusters contained cells from both L1-CRISPRi and control organoids, and we found no apparent difference in the contribution of L1-CRISPRi organoids to the different clusters, suggesting that L1s do not have a major influence on developmental fate in cerebral organoids (Figure 6f).

**Figure 6.**
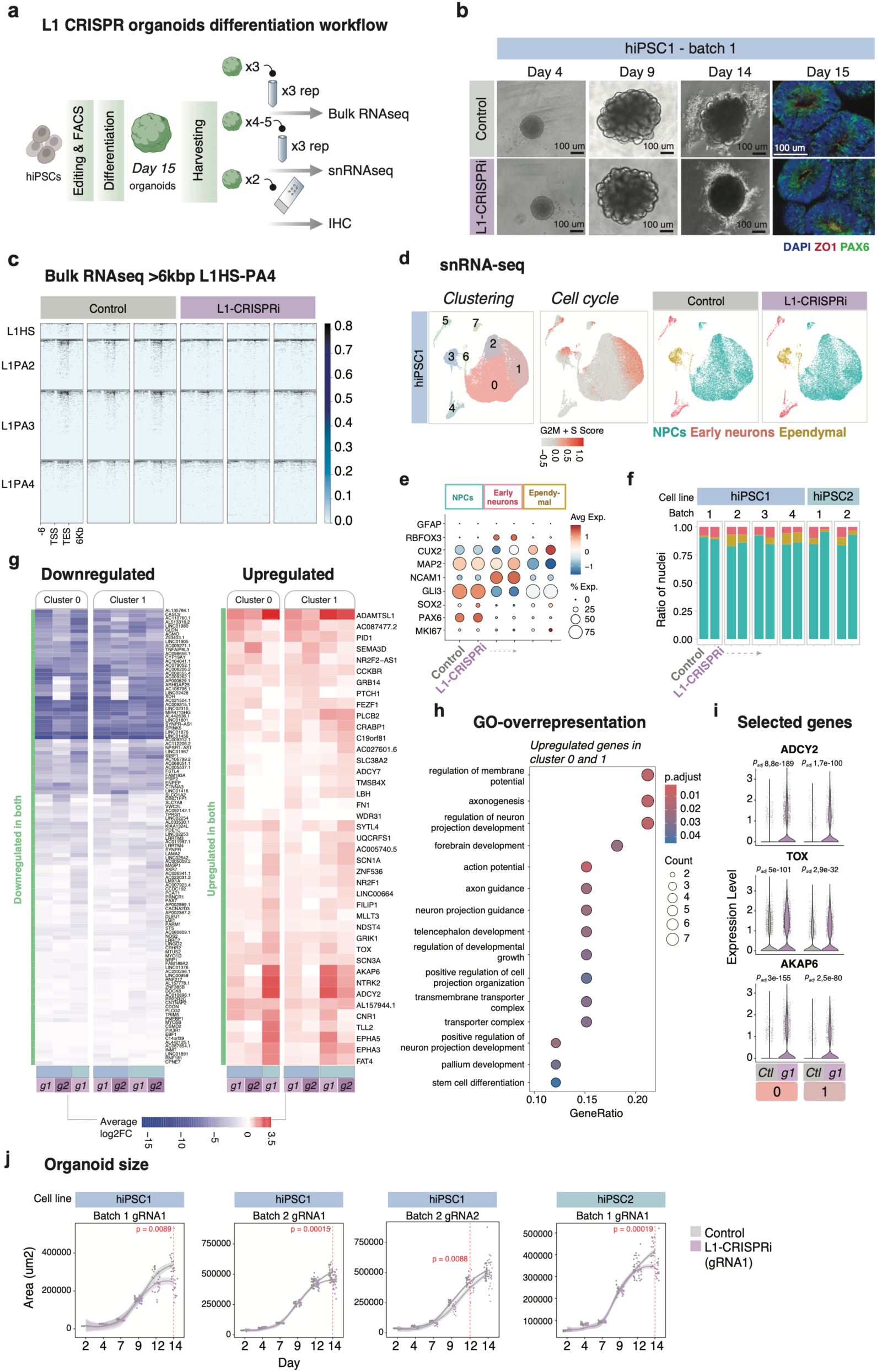
L1s regulate the exit of pluripotency and early neural differentiation in cerebral organoids. a) Workflow for the differentiation of L1-CRISPRi and control hiPSCs into day 15 unguided cerebral organoids and downstream analyses. b) Differentiation of the unguided cerebral organoids (hiPSC1 control vs. L1-CRISPRi). Brightfield images included from day 4, 9 and 14 of differentiation. Right: immunostaining showing neural rosettes and expression of the tight junction marker ZO1 (red) and the neural progenitor cell marker PAX6 (green). Nuclear staining DAPI is included in the overlay (blue). c) Normalised expression (RPKM) of uniquely mapped evolutionary young (L1HS-L1PA4), >6kbp L1s in control vs. L1-CRISPRi unguided cerebral organoids at day 15 of differentiation. d) Left: UMAP showing clustering of day 15 organoids (control + L1-CRISPRi). Middle: UMAP colored by cell cycle score (S + G2M). Right: UMAPs showing the identified cell types in control and L1-CRISPRi day 15 cerebral organoids. e) Dotplot showing the expression of selected markers used for the characterization of the cell clusters (dot size showing the percentage of cells expressing the gene, color indicates average expression in cell type per condition). f) Barplot of the cell types distribution across batches and cell lines of control vs. L1-CRISPRi organoids. g) Left: heatmap showing all the downregulated genes (L1-CRISPRi vs. control organoids) found in both clusters 0 and 1 across batches and cell lines (avg log_2_FC>1). Right: heatmap with all the upregulated genes found in clusters 0 and 1 across batches and cell lines (avg log2FC>1). h) Gene set enrichment analysis of upregulated genes from cluster 0 and 1 (L1-CRISPRi vs. control organoids). i) Violin plots of selected genes upregulated in L1-CRISPRi (purple) compared to control organoids (grey) in cluster 0 and cluster 1 (Seurat::FindMarkers, Wilcoxon test). j) Growth curves showing the size measurements distribution of control vs. L1-CRISPRi cerebral organoids from day 2 to day 15 of differentiation. Control is grey, L1-CRISPRi (gRNA1) purple. Area is measured as μm2 and was assessed with Fiji ImageJ. Student T-test was performed to compare the two conditions in each time point.

Next, we analyzed the transcriptional difference between L1-CRISPRi and control organoids. We first confirmed the transcriptional silencing of the L1s by using an in-house bioinformatics pipeline that allows for the analysis of L1 expression from the snRNA-seq dataset^26,49^. By backtracking the reads from the cells that make up each cluster, we are able to create a pseudo-bulk population of cells by cluster and analyse L1 expression using unique mapping. This pseudo-bulk approach greatly increases the sensitivity of L1 analysis and allows for a quantitative estimation of L1 expression at single cell type resolution. We found clear evidence of L1 expression from multiple loci in most clusters in the control organoids, as well as evidence of silenced L1 expression in the L1-CRISPRi organoids (Supplementary Figure 6c). In addition, we confirmed that several of the L1 *cis*-regulated genes (see Figure 4c) were also silenced in all clusters in the L1-CRISPRi organoids (Supplementary Figure 6d).

In the two largest NPC clusters (cluster 0 & cluster 1), representing non-proliferating and proliferating NPCs respectively, we found a set of genes that were consistently up- or downregulated in L1-CRISPRi organoids among the different batches, in both cell lines, and using either of the gRNAs (Figure 6g). Downregulated genes (n=116, log2FC < -0.25) were predominantly located in the close vicinity of an L1 and many of these genes were the same as the *cis*-regulated genes identified in the iPSCs (Figure 6g). Interestingly, many of the upregulated genes (n=41, log2FC > 0.25), which likely represent downstream consequences of the L1-silencing, have been linked to neuronal growth and neurogenesis, including *AKAP6, TOX, NTRK2* and *SCN3A^50–53^* (Figure 6g-i). Indeed, gene set enrichment analysis of the consistently upregulated genes in the major NPC clusters (cluster 0 and cluster 1) identified terms such as central nervous system differentiation, forebrain and synapse development, driven by genes related to functions in cAMP and calcium signaling such as *AKAP6* and *ADCY2* (Figure 6h-i)^50,54,55^. Thus, the transcriptome of NPCs in organoids that lack L1 expression suggests a more advanced differentiated profile when compared to control cerebral organoids, implying that L1 expression is important for an appropriate progress of the finely tuned neural differentiation. In line with this, we found that quantification of organoid size throughout differentiation revealed that L1-CRISPRi organoids were reproducibly and significantly smaller than control organoids. This difference appeared after 10 days and increased until day 15, the last time point quantified (Figure 6j). The results were reproduced in three independent batches using the two different cell lines and the two different gRNAs (Figure 6j, t-test hiPSC batch 1 p=0.0089, batch 2 p=0.00015, hiPSC2 p=0.00019, gRNA2 p=0.0088).

Together, these results show that silencing of L1s results in organoids containing the same cell types as control organoids, suggesting that L1s do not influence developmental fate. However, we found that the L1-CRISPRi organoids exhibited transcriptome changes consistent with alterations in neural development in NPCs, resulting in organoids that are reduced in size. These observations are consistent with an important role for L1 expression in the exit from pluripotency and early human brain development.

## Discussion

Several recent studies have shown that L1s are expressed in early mammalian development, where they are proposed to play an important regulatory role^11,12,38^. However, the abundance and repetitive nature of L1s make it difficult to accurately measure their transcription, particularly at individual loci^32^. Thus, the majority of these recent studies have largely considered L1s as a single family thereby limiting mechanistic analysis of how L1s may influence early developmental programs. In our study, we address this issue in human iPSCs and cerebral organoids using a detailed multiomics analysis, including long-read sequencing and custom bioinformatics pipelines. Our results show that thousands of unique L1 loci are expressed in human pluripotent stem cells and during early neural differentiation, where they add a hominoid-specific layer of transcriptional complexity.

We find that L1 expression in human iPSCs is largely restricted to human-specific (L1HS) and hominoid-specific (L1PA2 & L1PA3) elements. Expression of these elements correlates with lower levels of DNA methylation, suggesting that hominoid-specific L1s escape repressive epigenetic mechanisms that otherwise silence TEs in pluripotent stem cells^56^. This is consistent with a previous study that found that ancestral L1PA3 elements underwent structural sequence changes that allow them and subsequent younger L1 families to escape silencing by KRAB zinc finger proteins^57^, a large family of transcription factors that mediate transcriptional silencing of TEs in early development. However, upon differentiation into cerebral organoids, evolutionary young L1s were largely silenced by DNA methylation. The mechanism underlying this phenomenon is unknown, but may be related to other TE repression mechanisms, such as those associated with the HUSH complex^58–60^. Importantly though, our data highlight that many L1 elements are completely demethylated and highly expressed in cerebral organoids and therefore do not follow this general rule. Several mechanisms may underlie this discrepancy. For example, the L1 integration site is likely to be important, and the presence of highly active nearby gene promoters or other regulatory elements may influence the epigenetic status of L1s and their expression^10,26,47,61,62^. In addition, single nucleotide variants or small deletions in the regulatory regions of individual L1 loci could prevent the recruitment of silencing factors^9^. Crucially, our data underscore that when analyzing L1 expression, it is not sufficient to investigate expression at the family level but rather approaches that allow single locus resolution are needed.

Since the expression of L1s in hiPSCs correlates with the presence of H3K4me3 and low levels of DNA methylation at the promoter region, our data demonstrate that the L1 promoter is active in human iPSCs. This promoter activity provides a rich source of hominoid-specific transcripts. To investigate how L1s influence the transcriptome of hiPSCs, we optimised a CRISPRi system that silences the L1 promoter with high precision and efficiency. A combination of RNA-seq, mass spectrometry proteomics and CUT&RUN epigenomic analysis confirmed on-target silencing of virtually all active L1 promoters in hiPSCs, resulting in loss of L1 RNA and protein and inhibition of L1-mediated *cis*-acting effects. We found that about 100 genes in human iPSCs, a third of which are protein-coding, depend on the activity of the L1 promoter. For the protein coding genes, the L1 antisense promoter is used as an alternative promoter, leading to the expression of hominoid-specific alternative isoform of these conserved protein-coding genes. Thus, L1s are wired into the transcriptional network of human pluripotent stem cells.

Despite the global loss of L1 promoter activity, we found that L1-CRISPRi hiPSCs remained in a proliferative pluripotent state as demonstrated by RNA-seq, mass spectrometry proteomics and immunocytochemistry analyses. This is somewhat at odds with two previous studies in mouse and human pluripotent stem cells^11,38^, which suggested that L1s are essential for maintaining a pluripotent state. However, these studies used different technical approaches to silence L1s, relying on antisense oligonucleotides targeting ORF1 or ORF2. The key advantage of our CRISPRi approach is that by targeting the gRNA to the 5’ end of L1s, we limit the targeted loci to full-length L1s that are transcriptionally active. This increases specificity as most L1s are 5’ truncated as a consequence of an imperfect retrotransposition event. Such fragmented L1s are present in very high numbers throughout the human genome and are often part of other transcripts, making it difficult to avoid off-target effects when targeting more central parts of the L1 sequence. On the other hand, our CRISPRi approach is limited because it cannot distinguish between *cis-* and *trans*-acting mechanisms. Theoretically, L1s can mediate regulatory functions either through *cis*-acting mechanisms on nearby genes, as documented in the current study, or through *trans*-acting mechanisms, where L1-derived RNA or proteins carry out other regulatory mechanisms throughout the cell. Thus, the *cis*-acting mechanisms that we find at protein coding genes are likely to be complemented by *trans*-acting L1 functions, which we did not investigate further in the current study. However, recent studies by us and others suggest that such *trans*-acting L1 mechanisms are likely to be important both in pluripotent cells and during brain development^11,26–29,38^.

Several of the L1-regulated genes we documented in this study have been implicated in brain development, suggesting that L1s may play an important role in this process. Our data show that cerebral organoids in which L1 expression was silenced were smaller in size and had an altered transcriptome of NPCs compared to control counterparts. Our findings are reminiscent of previously observed differences when comparing human cerebral organoids with those derived from non-human great apes, implicating L1s in the evolution of the human brain^63–65^. In addition, many of the genes we found upregulated in L1 organoids have been implicated in neurodevelopmental and psychiatric disorders. As L1s are highly polymorphic within the human population, the prevalence of particular L1 copies, single nucleotide polymorphisms and structural variants in fixed L1s in the genome could therefore influence the etiology of brain disorders through altered brain development^66,67^.

In conclusion, our results illustrate how L1s provide a layer of hominoid-specific transcriptional complexity in human pluripotent stem cells that is important for the exit of pluripotency and early brain development. Thus, L1s represent a set of genetic material that contributes to important gene regulatory and transcriptional networks in early human development that could have contributed to human brain evolution. Our findings suggest that L1s should no longer be neglected and that these sequences must be included as individual loci in future investigations to study human evolution and the underlying genetic causes of human disorders.

## Methods

### Cell culture

Two hiPSC lines generated by mRNA transfection were used in the study (RBRC-HPS0328 606A1 and RBRC-HPS0360 648A1 [RIKEN]; referred to as hiPSC1 and hiPSC2, respectively). The cells were cultured as previously described^68,69^. Briefly, hiPSCs were maintained on Lam521-coated (0.7 μg/cm^2^ [Biolamina]) Nunc Δ multidishes in iPS media (StemMACS iPS-Brew XF and 0.5% penicillin/streptomycin [GIBCO]). The cells were passaged every 2-4 days until 70-90% confluency. At passaging, cells were rinsed once with DPBS (GIBCO) and dissociated with Accutase (250 μl/well in 12-wells plates) [GIBCO]) for 7-10 minutes at 37 °C. After incubation, the cells were collected, transferred to 10 ml of wash medium (9 ml DMEM/F-12 [GIBCO]; 1 ml Knockout Serum Replacement [GIBCO]), and centrifuged at 400 x g for 5 minutes. The cells were resuspended in 1 ml iPS media and plated with 10 μM of Y27632 (Rock inhibitor [Miltenyi]) for expansion. Media was changed daily.

### Bulk RNA sequencing

Total RNA was isolated from cells using the RNeasy Mini Kit (Qiagen). The sequencing libraries were then generated using Illumina TruSeq Stranded mRNA library prep kit (with poly-A selection) and sequenced on a NovaSeq6000 or Novaseq X plus (paired end 2 × 150bp). Detailed protocol can be found at DOI: dx.doi.org/10.17504/protocols.io.36wgqjqbkvk5/v1.

### Bulk RNA-seq analysis

#### TE subfamily quantification

For the quantification of TE subfamilies, the reads were mapped using STAR aligner (version 2.6.0c; RRID:SCR_004463)^70^ with an hg38 index and GENCODE version 38 as the guide GTF (--sjdbGTFfile), allowing for a maximum of 100 multimapping loci (--outFilterMultimapNmax 100) and 200 anchors (--winAnchorMultimapNmax). The rest of the parameters affecting the mapping was left in default as for version 2.6.0c.

The TE subfamily quantification was performed using TEcount from the TEToolkit (version 2.2.3; RRID:SCR_023208) in mode multi (--mode). GENCODE annotation v38 was used as the input gene GTF (--GTF), and the provided hg38 GTF file from the author’s web server was used as the TE GTF (--TE).

#### TE quantification

Reads were mapped using STAR aligner (version 2.6.0c; RRID:SCR_004463) with an hg38 index and GENCODE version 38 as the guide GTF (--sjdbGTFfile). We allowed a single mapping locus (--outFilterMultimapNmax 1) and a ratio of mismatches to the mapped length of 0.03 (--outFilterMismatchNoverLmax).

We measured in-sense and in-antisense transcription over features as previously described^26^ (see methods sections “Bulk RNAseq: TE quantification” and “Bulk RNAseq: Comparison between sense and antisense transcription over TEs”). For consistency (and to avoid quantifying over simple repeats, small RNAs, and low-complexity regions), we quantified TEs using the curated hg38 GTF file provided by the TEtranscripts authors. We considered an element to be “expressed” in a particular condition if at least one sample’s normalized expression was higher than two, i.e., TE counts divided by the sample distances (sizeFactor) as calculated by DESeq2 using the gene quantification (see the “Bulk RNA-seq analysis: Gene quantification” section).

To create in-sense and in-antisense deeptools heatmaps of evolutionary young L1 subfamilies, we used DeepTools’ (version 2.5.4; RRID:SCR_016366) computeMatrix, computeMatrixOperations, and plotHeatmap functions as previously described26 (the method section “Bulk RNAseq: Transcription over evolutionary young L1 elements in bulk datasets”).

#### Gene quantification

Using uniquely-mapped reads, genes were quantified using featureCounts from the subread package (version 1.6.3; RRID:SCR_012919) forcing strandness (-s 2) to quantify by gene_name (-g) from the GTF of GENCODE version 38.

#### Differential gene expression analysis

We performed differential expression analysis using DESeq2 (version 1.44.0; RRID:SCR_015687) with the read count matrix from featureCounts (subread version 1.6.3; RRID:SCR_012919) as input. Fold changes were shrunk using DESeq2:: lfcShrink.

For the produced heatmaps, counts were normalized by median of ratios as described by Love *et al.*71, summed with a pseudo-count of 1 and log^2^-transformed.

For further detail, please refer to the Rmarkdown on the GitHub.

#### Differential TE subfamilies expression analysis

We performed differential expression analysis using DESeq2 (version 1.44.0; RRID:SCR_015687) per guide RNA, per cell line, and per experiment (some experiments were reproduced on a second batch of organoids) using the uniquely-mapped read quantification of TEs (see section above “Bulk RNAseq: TE quantification”). Fold changes were shrunk using DESeq2:: lfcShrink.

Using the gene DESeq2 object (see section above), we normalized the TE subfamily counts by dividing the read count matrix by the sample distances (sizeFactor) as calculated by DESeq2 using the gene quantification (see the “Bulk RNA-seq analysis: Gene quantification” section). For heatmap visualization, a pseudo-count of 1 was added and log2-transformed.

### Western blot

RIPA buffer (Sigma-Aldrich) and a complete protease inhibitor cocktail were used to lyse the cells on ice for 30 minutes. The cells were then pelleted at 17 000xg, 4° C, for 20 minutes. Supernatants were collected and mixed with Bolt LDS loadig buffer 4x (Novex) and Bolt reducing agent 10x (Novex) and boiled at 95 °C for 5 minutes prior separation on a 4–12% Tris-glycine SDS-PAGE gel (200 V, 45 min). Proteins were then transferred from the gel to a PVDF membrane with the Transblot-Turbo Transfer system (BioRad). The membrane was blocked for 1 h in TBST with 5% skimmed milk (MTBST) before incubation overnight at 4° C with mouse anti-L1-ORF1p [Millipore MABC1152, 1:1,000 dilution] diluted in MTBST. The following day, the membrane was washed twice for 15 minutes in TBST and incubated for 1 h at room temperature with HRP-conjugated anti-mouse secondary antibody (Cell signalling, 1:5000) diluted in MTBST. After washing the membrane twice in TBST and once in TBS, the protein was detected by chemiluminescence using ECL Select reagents (Cytiva) according to manufacturer’s instructions and then imaged with a Chemi-Doc system (BioRad). Finally, the membrane was stripped using the Restore PLUS Western Blot Stripping Buffer (Thermo) as per instructions, re-blocked for 1 h in MTBST after which the procedure for the β-actin staining (HRP-472 conjugated anti-β-actin, Sigma A3854, 1:50 000 dilution). was performed as above. Detailed protocol can be found at DOI: dx.doi.org/10.17504/protocols.io.ewov1d562vr2/v1

### Immunocytochemistry

*HiPSCs staining.* Cells were rinsed once with DPBS and fixed for 15 minutes with 4% paraformaldehyde (Merck Millipore). The fixed cells were then rinsed three times with DPBS, and then pre-blocked for ∼1h in blocking solution (KPBS with 0.25% Triton X-100 [FisherScientific] and 5% normal donkey serum). The primary antibodies were added to the blocking solution and incubated overnight at 4 °C. On the following day, cells were washed twice with KPBS and the secondary antibodies, diluted in the blocking solution, were added and incubated at room temperature for 2 hours. Stained cells were rinsed once with KPBS, then stained with DAPI (1:1000 dilution, [Sigma-Aldrich]) for 10 minutes. Finally, 2 rinses with KPBS were performed before imaging. Stainings were imaged on a Leica microscope (model DMI60000 B), and images were cropped and adjusted on Fiji ImageJ.

Antibodies used: mouse anti-OCT3/4 (1:300 dilution [Santa Cruz, RRID: AB_628051]); rabbit anti-NANOG (1:600 dilution [Abcam, RRID: AB_446437]); mouse anti-hORF1p, clone 4H1 (1:300 dilution [Sigma-Aldrich, RRID: RRID:AB_2941775]); donkey anti-mouse Cy3 (1:550 dilution [Jackson Lab]); donkey anti-rabbit Alexa647 (1:550 dilution [Jackson Lab]).

The protocol can be found at DOI: dx.doi.org/10.17504/protocols.io.5qpvor7pdv4o/v1

### CUT&RUN

Two different CUT&RUN protocols were used depending on the target. A standard CUT&RUN protocol was applied to profile H3K4me3 and H3K9me3. A crosslinking step was added to profile dCas9 in L1 CRISPRi hiPSCs.

#### Standard CUT&RUN

The protocol performed follows a previously described CUT&RUN protocol^34,72^. Briefly, 500k cells were harvested, washed twice in Wash buffer (20 mM HEPES pH 7.5, 150 mM NaCl, 0.5 mM spermidine, 1 Roche Complete Protease Inhibitor EDTA-free tablet diluted in dH^2^O) and then attached to 10 ul of pre-activated ConcavalinA-coated magnetic beads. 10 ul of beads per sample were pre-activated in Binding buffer (20 mM HEPES pH 7.9, 10 mM KCl, 1 mM CaCl_2_, 1 mM MnCl_2_) and kept on ice until use. The bead-bound cells were then placed in a magnetic stand and, after carefully removing the supernatant, resuspended in ice-cold Antibody buffer (20 mM HEPES pH 7.5, 0.15 M NaCl, 0.5 mM spermidine, 1 Roche Complete Protease Inhibitor EDTA-free tablet, 0.05% w/v digitonin, 2 mM EDTA, 1:100 primary antibodies). The cells were left incubating in Antibody buffer overnight at 4 °C on gentle shake. On the following day, samples were placed on the magnetic stand, the liquid removed, and the beads were washed twice with Digitonin buffer (Wash buffer + 0.05% w/v digitonin). 50 ul of Protein A-MNase at a concentration of 700 ng/ml diluted in digitonin buffer were then carefully added to the beads while gently vortexing. The samples were incubated with the Protein A-MNase for 1h at 4 °C on gentle shake. Bead-bound cells were washed twice with 1 ml of Digitonin buffer (samples placed in the magnetic stand, liquid removed without dislodging the beads) and then resuspended in 100 ul of Digitonin buffer and cooled down to 0-2 °C for 5 minutes. Genome digestion was achieved by adding 2 mM CaCl_2_ to the samples which were then kept at 0 °C for 30 minutes. The reaction was quenched with 100 ul of 2X Stop buffer (0.35 M NaCl, 20 mM EDTA, 4 mM EGTA, 0.05% digitonin, 50 ng/ml of RNAse A, 50 ng/ml of glycogen, 10 ng/ml spike in DNA, all diluted in distilled H_2_o) and vortexing. After a 30 minutes incubation at 37 °C to release the DNA fragments, samples were placed on the magnetic stand and the supernatant was collected into a new 1.5 ml Eppendorf tube for DNA extraction via spin-column (NucleoSpin clean-up kit [Macherey-Bagel]).

Antibodies used with this protocol: anti-H3K9me3 (1:100 dilution [Abcam]); anti-H3K4me3 (1:100 dilution [Abcam]); IgG (1:100 dilution [Abcam]).

Detailed protocol can be found at DOI: dx.doi.org/10.17504/protocols.io.36wgqdb83vk5/v1

#### Crosslinked CUT&RUN

The protocol was the same as above except for some additional reagents in the buffers and the cross-linking (and cross-linking reversion) steps. Briefly, cells were centrifuged at 600xg for 3 minutes and the pellet was then resuspended in 1 ml of 0.1% formaldehyde solution diluted in the cell culture media to allow light crosslinking. After 1 minute incubation at room temperature, glycine was added to a final concentration of 125 mM, and the cells were centrifuged again at 600xg for 3 minutes. The supernatant was removed, and the cells resuspended in 800 ul of XL-Wash Buffer (20 mM HEPES pH 7.5, 150 mM NaCl, 0.5 mM spermidine, 1 Roche Complete Protease Inhibitor EDTA-free tablet, 1% Triton X-100, 0.05% SDS, diluted in dH_2_O). Cells were centrifuged at 1000xg for 3 minutes and resuspended again in XL-Wash Buffer. Centrifugation was performed at 1200xg for 3 minutes two times more, and the pellet resuspended in XL-Wash Buffer. 10 ul of pre-activated ConA-coated beads was added per sample and allowed to bind (5-10 minutes incubation on an end-over-end rotator). After placing the samples on the magnetic stand, cleared supernatant was removed and beads were resuspended in 100 ul XL-Antibody Buffer (XL-Wash Buffer, 0.05% digitonin, 2 mM EDTA, 1:50 anti-Cas9 antibody), in which they were incubated overnight at 4 °C on gentle shake. On the following day, samples were placed on a magnetic stand and, after removing the supernatant, resuspended in XL-Digitonin buffer (XL-Wash buffer, 0.05% digitonin). The wash was repeated once more and then the beads were resuspended in 50 ul/sample of Protein A-MNase diluted in XL-Digitonin buffer (final concentration 700 ng/ml), followed by 1 hour incubation at 4 °C on gentle shake. Bead-bound samples were washed then twice with XL-Digitonin buffer and then resuspended in 100 ul of XL-Digitonin buffer and cooled down to 0-2 °C for 5 minutes. 2ul of 100 mM CaCl2 was added to each sample and then incubated at 0°C for 30 minutes. The reaction was stopped by adding 2X Stop buffer (see: Standard CUT&RUN protocol) and incubated at 37°C for 30 minutes to release the fragmented chromatin. The samples were placed on the magnetic stand and the supernatant was collected in a new Eppendorf tube, where 0.09% SDS and 0.22 mg/ml of Proteinase K were added to reverse the crosslinking and incubated overnight at 55°C. DNA was extracted using NucleoSpin PCR Cleanup columns (Macherey-Nagel).

The antibody used in this step is anti-Cas9 (1:50 dilution [Takara]).

A detailed protocol can be found at DOI: dx.doi.org/10.17504/protocols.io.bp2l6dr5rvqe/v1

### CUT&RUN analysis

Paired-end reads (2 × 75) were aligned to the human genome (hg38) using bowtie2 (version 2.4.4; RRID:SCR_016368)^73^ (–local –very-sensitive-local –no-mixed –no-unal –no-discordant –phred33 -I 10 -X 700), converted to BAM files with SAMtools (version 1.9; RRID:SCR_002105) and sorted (SAMtools version 1.9; RRID:SCR_002105). Reads per kilobase per million mapped reads (RPKM) normalized bigwig coverage tracks were made with bamCoverage (DeepTools, version 2.5.4; RRID:SCR_016366)^74^. To normalize the signal from the dCas9 CUT&RUN bigwig files, we performed a subtraction (--ratio) using bigwigCompare of dCas9 gRNA1 cells signal minus the dCas9 control cells signal.

Tag directories were created using Homer (version 5.1; RRID:SCR_010881)^75^ makeTagDirectory on default parameters. Peak calling was performed using findPeaks (Homer), using the option “factor” as style for dCas9 CUT&RUN (-style). The rest of the parameters were left on default options. A window of 100 bp was added at both sides of the significant peaks (p-value < 0.05, as reported by homer) and only dCas9 L1-CRISPRi peaks that had no overlap in dCas9 control peaks were kept as true gRNA binding sites (bedr::in.region). and intersected (BEDtools, version 2.30.0; RRID:SCR_006646) to bed files containing coordinates of >6-kbp L1HS or any other L1PA subfamily (on-target; bedtools intersect; repeatmasker hg38 version open-4.0.5). The rest (-v) of the called peaks were considered off-targets. Matrices for heatmaps (both on- and off-target ones) were created using computeMatrix (DeepTools, version 2.5.4; RRID:SCR_016366) and visualized using plotHeatmap (DeepTools).

Tag directories of histone marks were created using Homer (version 5.1; RRID:SCR_010881)^75^ makeTagDirectory on default parameters. Peak calling was performed using findPeaks (Homer, version 5.1), using the option “histone” as style for the H3K4me3 and H3K9me3 CUT&RUN (-style). To identify active promoters among full-length evolutionary young L1s, the significant peaks (p-value < 0.05, as reported by homer) were intersected (BEDtools, version 2.30.0; RRID:SCR_006646) to a bed file containing the coordinates of the first 900bps of >6-kbp L1HS, L1PA2, L1PA3 and L1PA4 (repeatmasker hg38 version open-4.0.5).

### Oxford Nanopore DNA sequencing

HMW DNA was extracted from frozen hiPSC pellets (500 000-1 million cells) and day 15 cerebral organoids (5 organoids) using the Nanobind HMW DNA Extraction kit (PacBio) following the manufacturer’s instructions. HMW DNA was eluted in 100 μL of Buffer EB. Size selection to enrich for fragments >5 kb was performed using the Short Read Eliminator XS kit (PacBio) according to the manufacturer’s instructions. HMW DNA quantity and quality were assessed by Nanodrop and Qubit from the top, middle, and bottom of each tube, and on a TapeStation 4200 (Agilent) using the Agilent Genomic DNA Screen Tape Assay (Ref No. G2991-90040, Edition 08/2015). HMW DNA was sheared to approximately 10kb fragment length using a g-TUBE (Covaris) following the manufacturer’s protocol. Library preparation for whole genome sequencing was performed using the SQK-LSK114 Ligation Sequencing kit (Oxford Nanopore Technologies) with >1 μg DNA input. Libraries were each sequenced seperately using a FLO-PRO114M PromethION Flow Cell R10.4.1 on a PromethION platform (Oxford Nanopore Technologies) at SciLifeLab Uppsala and *in-house* on a PromethION 2 Solo (Oxford Nanopore Technologies) for 72 hours. Raw sequencing data (POD5 format) were basecalled and mapped to the human reference genome (GRCh38/hg38) using Dorado version 0.5.1-CUDA-11.7.0 (https://github.com/nanoporetech/dorado) utilizing the super accurate basecalling model <u>dna_r10.4.1_e8.2_400bps_sup@v4.3.0</u> and <u>dna_r10.4.1_e8.2_400bps_sup@v4.3.0_5mCG_5hmCG@v1</u> for methylation aware basecalling. Resulting BAM files were sorted and indexed using SAMtools version 1.18^76^.

Locus-specific L1 and L1 promoter (first 900 bp) methylation status were visualized using MethylArtist version 1.2.677 using the segmeth (--motif CG, with hg38 as reference genome) and locus functions with default parameters.

### Long-read direct RNA sample preparation, sequencing and analysis

#### RNA extraction

RNA was extracted from one hiPS cell pellet (3 million cells) using the Trizol RNA extraction protocol (Trizol reagent [Invitrogen #15596026]). Briefly, 1 ml of Trizol was added to the pellet to lyse the cells. The lysate was incubated for 5 minutes at room temperature (RT), and then 200 µl of chloroform were added followed by thorough mixing. The sample was then incubated at RT for an additional 3 minutes prior centrifugation at 12000 x g at 4 °C for 20 minutes. The aqueous phase (containing RNA) was transferred to a new tube where 500 µl of isopropanol where added prior a 10 minutes incubation at 4 °C. After that, RNA was pelleted via centrifugation at 12000 x g at 4 °C for 10 minutes. The RNA was resuspended in 1 ml of freshly prepared 75% ethanol, vortexed, and centrifuged for 5 minutes at 7500 x g at 4 °C. The pelleted RNA was air-dried and finally resuspended in 35 µl of RNase-free water. Lastly, the suspension was incubated at 56 °C for 15 minutes.

#### Poly(A) mRNA isolation

The NEBNext High Input Poly(A) mRNA Isolation Module (NEB #E3370S) protocol was used to select the mRNA for long-read direct RNA sequencing. Shortly, 20 µl of High Input Oligo d(T)_25_ Beads were washed twice in 100 µl of RNA Binding Buffer (2X). The supernatant was removed by placing the beads in a magnetic rack until the solution was clear. 50 µl of RNA Binding Buffer (2) were then added to the isolated RNA (see: *RNA extraction*), which was previously diluted to 50 µl. The RNA was denaturated by heating the sample at 65 °C for 5 minutes and then cooling it to 4 °C. Beads were then resuspended and incubated at room temperature for 5 minutes prior to placing them on the magnetic holder and removing the supernatant. Two additional washes were then performed with 200 µl of Wash Buffer. After removing the Wash Buffer, beads were thoroughly resuspended in 50 µl of Tris Buffer and then heated to 80 °C for 2 minutes, followed by a cool down to 25 °C. A second purification step followed with the addition of 50 µl of RNA Binding Buffer to increase the specificity of mRNA binding. The mRNA-bound beads were again washed with 200 µl of Wash Buffer. The mRNA was then finally eluted with 17 µl of Tris Buffer and incubation at 80 °C for 2 minutes, followed by cool down to 25 °C. The sample was replaced on the magnetic rack, and 15 µl of eluted RNA were transferred to a clean nuclease-free PCR tube and placed on ice immediately. The yield of the purified mRNA was assessed using a Qubit 3 fluorometer and Qubit RNA Assay kit.

#### ONT direct RNA sequencing library preparation and data analysis

The library for long-read direct RNA sequencing was prepared according to manufacturer’s protocol and using the Direct RNA Sequencing Kit (SQK-RNA004) from Oxford Nanopore Technologies. The sample was sequenced using a PromethION 2 Solo IT on a PromethION RNA Flow Cell (FLO-PRO004RA). Data was basecalled using dorado (version 0.7.1) with model rna004_130bps_sup@v3.0.1 and --modified-bases-models using dorado’s model rna004_130bps_sup@v3.0.1_m6A_GRACH@v1. Reads were mapped using minimap2 (-ax splice -uf -k14 -y) using hg38 as the reference genome. Visualization was done using the sorted and indexed bam file (samtools 1.16.1) on IgV (version 2.18.2).

### CRISPR inhibition (CRISPRi)

To silence the transcription of LINE-1s in hiPSCs, we adapted the protocol detailed previously^35,78^. The single guide sequences designed to recognize the 5’ UTR of full-length young L1 elements were previously published^36^. The guide sequences were inserted into a deadCas9-KRAB-T2A-GFP lentiviral backbone, pLV hU6-sgRNA hUbC-dCas9-KRAB-T2a-GFP, a gift from Charles Gersbach (Addgene plasmid #71237 RRID: Addgene_71237), using annealed oligos and the BsmBI cloning site. Lentivirus was produced as described below (see: Lentiviral production). hiPSCs were transduced with MOI 10 of LacZ and LINE-1-targeting gRNA lentiviral particles. Guide efficiency was validated using bulk RNA sequencing, due to the repetitive nature of the targets.

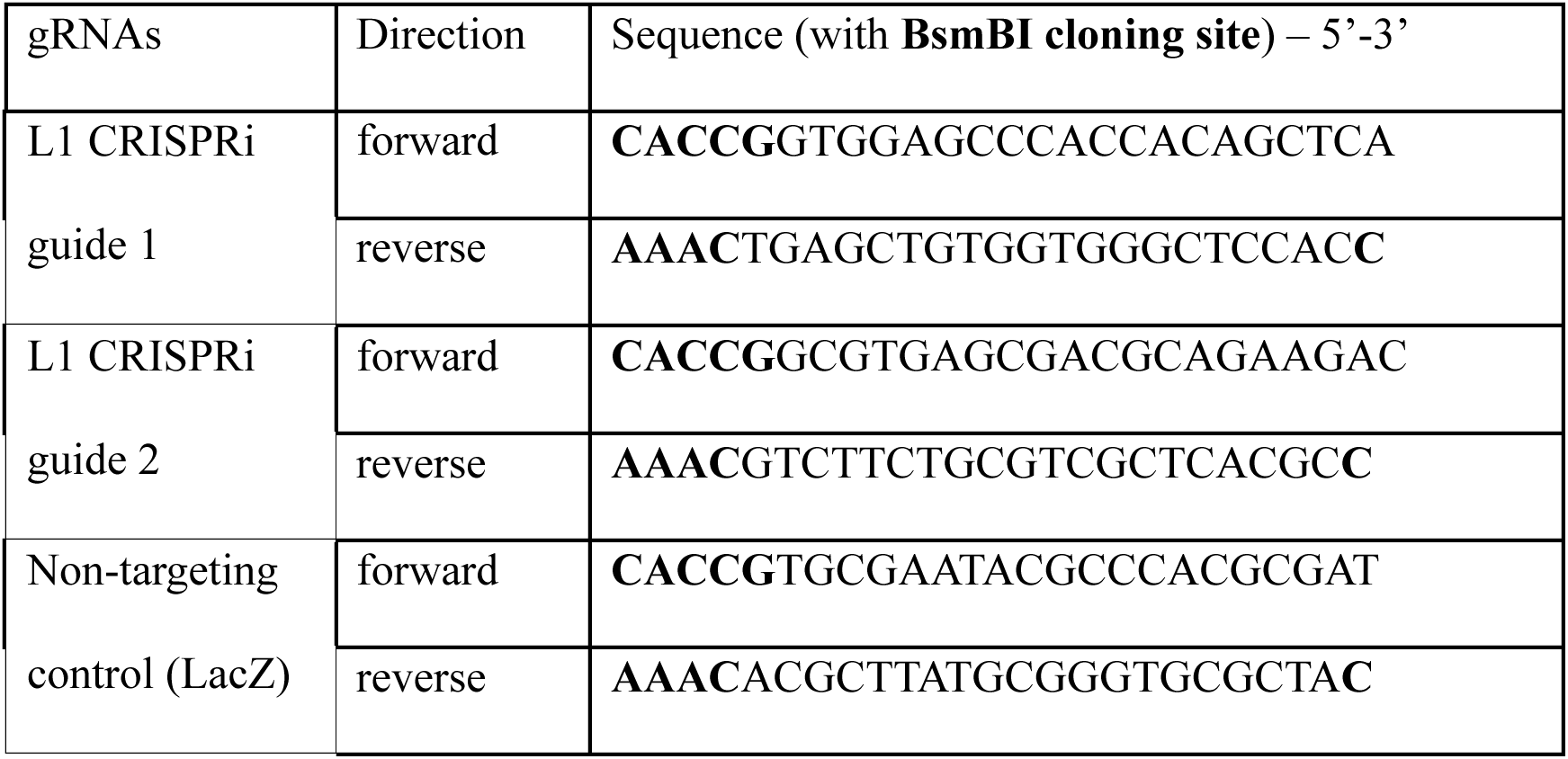

Between 7 and 10 days after transduction, GFP-positive cells were selected via FACS (FACS Aria, BD Biosciences). Briefly, cells were detached as described above (see: *Cell culture*), then resuspended in iPS media containing RY27632 (10 μM) and Draq7 (1:1000, BD Bioscience), and strained with a 70μm (BD Bioscience) filter. Gating parameters were determined by side and forward scatter to eliminate debris and dead cells. The GFP-positive gates were set using untransduced hiPSCs. Sorting gates and strategies were validated via reanalysis of sorted cells with a requirement of > 95% purity cut-off. 100 000-200 000 GFP-positive/Draq7-negative cells were collected for each sample, spun down at 400 x g for 5 min and snap-frozen on dry ice (for bulk RNA sequencing). Cell pellets were kept at −80°C until RNA was isolated. 300000 GFP-positive/Draq7-negative cells per samples were also collected, spun down at 400 x g for 5 min, resuspended in iPS media containing RY27632 (10 μM) and expanded as described above (see: Cell culture). GFP-positive cells were also frozen down for further use (e.g. cerebral organoids differentiation).

Detailed protocol can be found at DOI: dx.doi.org/10.17504/protocols.io.4r3l29ezqv1y/v1

### Lentiviral production

Lentiviral vectors were produced according to Zufferey *et al* ^79^. and were titered by qRT-PCR. Shortly, HEK293T cells (RRID: CVCL_0063) were grown to a confluency of 70 – 90% for lentiviral production and co-transfected with third-generation packaging and envelope vectors (pMDL (Addgene #12251), psRev (Addgene #12253), pMD2G (Addgene #12259)) together with Polyethyleneimine (PEI Polysciences PN 23966) in DPBS (GIBCO) in conjunction with the previously generated plasmids (see above: CRISPR inhibition). The lentiviruses were then harvested two days after transfection. The media was collected, filtered and centrifuged at 19500 x g for 2 hours at 4° C. The supernatant was removed from the tubes, the pellet resuspended in cold DPBS and left at 4° C overnight. The resulting lentivirus was aliquoted, titered, and stored at −80°C.

Detailed protocol can be found at DOI: dx.doi.org/10.17504/protocols.io.8epv5ro85g1b/v1

### Mass spectrometry data generation and analysis

#### Sample preparation for cellular proteome analysis

Pellets corresponding to 250,000 cells from L1 CRISPRi hiPSCs samples and the LacZ control (3 replicates per condition) were solubilized in 50 mM ammonium bicarbonate (Sigma) with 0.1% RapiGest (Waters) to denature proteins and shaken on a thermomixer (Eppendorf) at 400 rpm for 15 minutes decreasing the temperature from 80°C to 56°C. This was followed by reduction of disulfide bonds with 0.1M dithiothreitol at 56°C, cysteine alkylation with 0.2 M iodoacetamide at room temperature in the dark, and digestion overnight at 37°C with sequencing grade modified trypsin (enzyme: protein ratio 1:50, Promega). Digested peptides were acidified with 10% trifluoroacetic acid (TFA) and RapiGest was precipitated by incubation at 37°C. Peptides were desalted with SepPak C18 cartridges (Waters), dried by vacuum centrifugation, and stored at −80°C until LC-MS analysis. To generate a spectral library for the deep proteomic profile for the L1 CRISPRi hiPSCs proteome, 15-20 μg of peptides pooled from each sample were fractionated into 8 fractions by high-pH reversed-phase (HpH-RP) fractionation. Fractioned samples were dried by vacuum centrifugation and stored at −80°C until LC-MS analysis.

#### Liquid chromatography and mass spectrometry (LC-MS)

LC-MS analyses were carried out on an Orbitrap Exploris 480 MS instrument with a reverse phase UltiMate 3000 UHPLC system via an EASY-Spray ion source equipped with FAIMS Pro (all Thermo Fisher Scientific). Peptides for the single-shot samples were measured with data-independent acquisition (DIA). Digested peptides were loaded onto a trap cartridge (Acclaim PepMap C18, 5 mm particle size, 0.3 mm inner diameter x 5 mm length, Thermo Fisher Scientific) and separated by EASY-Spray analytical column (2 mm particle size, 75 mm inner diameter x 500 mm length, Thermo Fisher Scientific). Each sample was injected twice and eluted with a linear gradient ranging from 2-19% Solvent B (0.1% formic acid (FA) in 80% acetonitrile (ACN) over 80 min, 19-41% B over 40 min, 41-90% B over 5 min and held at 90-95% B for 5 min at a constant flow rate of 300 nl/min at 45°C. For data-dependent acquisition (DDA) and DIA measurements, the spray voltage was set at 2.1 kV, ion transfer tube temperature was set at 2750C, and FAIMS compensation voltages (CV) were set to -45 and -60. For DIA, peptides were analyzed with one full scan (350–1,400 m/z, R = 120,000) at a normalized AGC target of 300%, followed by 38 DIA MS/MS scans (350–1,050 m/z) in HCD mode (isolation window 18 m/z, 1 m/z window overlap, normalized collision energy 27%), with fragments detected in the Orbitrap (R = 15,000). The fractions were measured with DDA to generate the spectral library. Each fraction was injected twice and eluted over 180 min gradients (UltiMate 3000 UHPLC, Thermo Fisher Scientific) ranging from 2-25% Solvent B (0.1%FA in 80% ACN) over 100 min, 25-40% B over 20 min, 40-90% B over 2 min and held at 90% B for 5 min at a constant flow rate of 300 nl/min at 45°C. The full MS scan was performed in the Orbitrap in the range of 350 to 1400 m/z at a resolution of 60,000 with automatic gain control (AGC) set to 300 and maximum ion injection time set to auto. The intensity threshold was set to 5.0e3 and mass tolerance to 10 ppm. The most intense ions selected in the first MS scan were isolated for higher-energy collision-induced dissociation (HCD) at a precursor isolation window width of 1.6 m/z, and normalized AGC of 75%. Maximum ion injection time for MS^2^ was set to auto with MS^2^ resolution set to 15,000. The first mass and the normalized collision energy were set to 110 m/z and 30%, respectively. All data were acquired in positive polarity and MS/MS for DIA and DDA were acquired in centroid and profile mode, respectively.

#### MS raw data processing and Statistical analysis

Spectronaut (version 18, Biognosys AG) was used to build the spectral library and analyze the single-shot DIA runs. For the generation of the hybrid library, the reverse phase fraction runs (DDA) were combined with the single-shot individual sample runs (DIA) and searched with ‘Pulsar’ using default settings. FDR was set to 1% to determine the significance level. The single-shot DIA samples were searched against the hybrid library with the human SwissProt reference proteome (20,602 entries downloaded on October 2024) and commonly used contaminants. Searches used carbamidomethylation as fixed modifications, methionine oxidation, and protein N-terminal acetylation as variable modifications. The Trypsin/P proteolytic cleavage rule was used, permitting a maximum of 2 missed cleavages and a minimum peptide length of 7 amino acids. ‘Cross run normalization’ was enabled with Normalization Strategy set to ‘local normalization’ based on rows with ‘Identified in All Runs (Complete)’. Data filtering was set to Q-value and the Q-value thresholds were set to 0.01 at PSM, peptide, and protein levels. Protein quantification and statistical analysis were performed with Msstats^80^ (version 4.12.0). Contaminants were filtered and features were converted to MSStats format for downstream processing. Uninformative features were removed, and missing values were imputed with the ‘MBimpute’ function within MSStats. For statistical analysis, MSStats ‘group comparison’ was performed for the gRNA samples against the LacZ control. Differentially expressed proteins were selected with adjusted p-value of less than 0.05 and a fold change of more than 1.5 for each comparison.

### Unguided cerebral organoids culture

Organoids were cultured using the STEMdiff^TM^ Cerebral Organoids kit and following the manufacturer’s protocol until day 15 of differentiation. Shortly, at day 0 of differentiation hiPSCs at 70-90% confluency were detached and centrifuged as described above (see Cell culture: *hiPSCs*), then resuspended in 1 ml of EB Seeding Medium (STEMdiff^TM^ Cerebral Organoids Basal Medium 1 and STEMdiff^TM^ Cerebral Organoids Supplement A, supplemented with 10 um of Y-27632 (Rock Inhibitor)). Cells were counted and the required volume to obtain 90000 cells/ml was calculated. 100 ul per well were then distributed in a round-bottom ultra-low attachment 96-well plate (9000 cells/well). Plate was left incubating undisturbed for 24 h. At day 2 and 4 of differentiation 100 ul of fresh EB Formation Medium (STEMdiff^TM^ Cerebral Organoids Basal Medium 1 and STEMdiff^TM^ Cerebral Organoids Supplement A, 4:1) were added to each well. On day 5, organoids were transferred to an ultra-low attachment 24-well plate with 0.5 ml of Induction Medium (STEMdiff^TM^ Cerebral Organoids Basal Medium 1 and STEMdiff^TM^ Cerebral Organoids Supplement B), in each well (1 organoids/well to avoid organoids fusion). Organoids were incubated at 37 °C for 48 h. On day 7, organoids were embedded with ∼15 ul of Matrigel per organoid and transferred to an ultra-low attachment 6- well plate with 3 ml of Expansion Medium (STEMdiff^TM^ Cerebral Organoids Basal Medium 2, STEMdiff^TM^ Cerebral Organoids Supplement C, STEMdiff^TM^ Cerebral Organoids Supplement D) per well. 10-12 organoids per well were cultured together. On day 10 and 13 media was changed with 3 ml/well of Maturation Medium (STEMdiff^TM^ Cerebral Organoids Basal Medium 2, STEMdiff^TM^ Cerebral Organoids Supplement E). On day 15 of differentiation, the organoids were collected for downstream analyses. 3 organoids were collected and snap-frozen for each bulk RNA sequencing replicate (n= 3 replicates per condition). 4-5 organoids were collected and snap-frozen for each snRNA seq replicate (n= 3 replicates per condition). 2 organoids were collected for immunostaining (see: Immunohistochemistry for samples preparation). 5 organoids were collected for long-read DNA sequencing.

### Immunohistochemistry

Organoids collected for staining were transferred into a 24-well plate and fixed with 4% paraformaldehyde (PFA) for 2 hours. Fixed organoids were then rinsed twice in DPBS and immersed in a 1:1 OCT (catalog no. 45830, HistoLab) and 30% sucrose emulsion in which they were incubated overnight at 4 °C on gentle shake. On the following day, the organoids were transferred to a cryomold in OCT and frozen on dry ice, to be stored at -80 °C until cryosectioning. Organoids were cut on a cryostat at -20 °C into sections of 20 um thickness and placed onto Superfrost Plus microscope slides.

A hydrophobic PAP Pen was used to draw the borders of the slides. Slides were then washed twice in 1X KPBS (PBS + 0.3% Triton X-100) and fixed again with 4% PFA for 10 minutes. After two washes with KPBS, slides were immersed in Blocking solution (TKPBS + 5% Normal Donkey Serum) for at least 1 h. ∼300 ul of Primary antibody solution (blocking solution + primary antibodies at the chosen dilution) were added to each slide and these were incubated overnight at 4 °C in a humidified chamber. On the following day, slides were washed three times with KPBS and then incubated for at least 1 h with ∼300 ul of Secondary antibody solution (blocking solution + primary antibodies at the chosen dilution, 1:1000 DAPI) in the dark. Slides were then washed three times with KPBS and finally coverslipped with FluorSave mounting medium. Slides were imaged on a confocal Microscope Leica TSC SP8 and images cropped and adjusted on ImageJ Fiji.

Antibodies used: mouse anti-ZO1(Invitrogen, 339100; 1:300); rabbit anti-PAX6 (Biolegend, 901301; 1:300); anti-mouse Cy3 (Jackson Lab; 1:550); anti-rabbit Alexa Fluor647 (Jackson Lab; 1:550).

### Single nuclei isolation

The nuclei isolation from organoids was performed as previously published^49,81^ using a sucrose gradient-based isolation. In brief, the organoids were thawed and dissociated in ice-cold lysis buffer (0.32 M sucrose, 5 mM CaCl_2_, 3 mM MgAc, 0.1 mM Na_2_EDTA, 10 mM Tris-HCl, pH 8.0, 1 mM DTT) with a 1 ml tissue douncer (Wheaton). The lysate was then carefully layered on top of a sucrose cushion (1.8 M sucrose, 3 mM MgAc, 10 mM Tris-HCl pH 8.0, and 1 mM DTT diluted in milliQ water) before centrifugation at 30,000 × g for 2 hours and 15 min. Once the supernatant was removed, the pelleted nuclei were softened for 10 min in 100 μl of nuclear storage buffer (15% sucrose, 10 mM Tris-HCl pH 7.2, 70 mM KCl, and 2 mM MgCl_2_, all diluted in milliQ water), resuspended in 300-800 μl of dilution buffer (10 mM Tris-HCl pH 7.2, 70 mM KCl, and 2 mM MgCl_2_, diluted in milliQ water) and then filtered through a cell strainer (70 μm). The nuclei were sorted via FANS (with a FACS Aria, BD Biosciences) at 4° C at low flow rate using a 100 μm nozzle (reanalysis showed >95% purity).

Detailed protocol can be found at DOI: dx.doi.org/10.17504/protocols.io.5jyl8j678g2w/v1.

### Single nuclei sequencing

The nuclei for single nuclei RNA sequencing (8500-10000 nuclei per sample) were loaded onto the Chromium Next GEM Chip G Single Cell Kit along with the reverse transcription mastermix according to manufacturer’s protocol for the Chromium Next GEM single cell 3’ kit (10X Genomics, PN-1000268) to generate single-cell gel beads in emulsion. cDNA amplification was done following the guidelines from 10X Genomics using 13 cycles of amplification of the 3’ libraries. Sequencing libraries were generated with unique dual indices (TT set A) and pooled for sequencing on a Novaseq6000 or Novaseq X plus using a 100-cycle kit and 28-10-10-90 reads.

### Single nuclei RNAseq analysis

#### Gene quantification

The raw base calls were demultiplexed and converted to sample-specific fastq files using 10x Genomics Cell Ranger mkfastq (version 6.0.0; RRID:SCR_017344)^82^. Cell Ranger count was run with default settings, using an mRNA reference for single-cell samples and a pre-mRNA reference (generated using 10x Genomics Cell Ranger 6.0.0 guidelines) for single-nucleus samples.

#### Clustering

Samples were analyzed using Seurat (version 5.1.0; RRID:SCR_007322)^83^. For each sample, cells were filtered out if the percentage of mitochondrial content was over 10% (perc_mitochondrial). Cells were discarded if the number of detected features (nFeature_RNA) was higher than two SDs over the mean in the sample or lower than a SD below the mean in the sample (to avoid low quality cells). Counts were normalized using LogNormalize method (Seurat::NormalizeData). Clusters were defined with a resolution of 0.1 (Seurat::FindClusters).

#### Gene differential expression analysis

We used Seurat’s FindMarkers grouped by cell types and on default parameters as for version 5.1.0 to identify differentially expressed genes (Wilcoxon test). A gene was considered to be differentially expressed on a cell type if its adjusted *P* value was below 0.05. To define the lists of genes consistently up or downregulated in all experiments (Figure 6f), we performed individual comparisons between L1-CRISPR and LacZ conditions in each cell type and guide. We used hiPS6 gRNA1 as our “base” comparison. If a gene had a log2FC > 0.25 in our “base” comparison and log2FC > 0 in the rest of the comparisons (hiPS6 gRNA1 vs LacZ, hiPS48 gRNA1 vs LacZ, and hiPS48 gRNA2 vs LacZ), the gene is considered to be a consistently upregulated gene. Similarly, if a gene had a log2FC < -0.25 in our “base” comparison and log2FC < 0 in the rest of the comparisons, the gene is considered to be a consistently downregulated gene.

#### TE quantification

We used an in-house pseudo-bulk approach to processing snRNA-seq data to quantify TE expression per cluster, similar to what has been previously described^49^. All clustering, normalization and merging of samples were performed using the contained scripts of get_clusters.R [get_custers() from the Sample class] and merge_samples.R [merge_samples() from the Experiment class] of trusTEr (version 0.1.2; DOI:10.5281/zenodo.14362613) using the unique mapping mode when processing clusters. Documentation of the pipeline can be found at https://molecular-neurogenetics.github.io/truster/.

Dependencies were ran using the following software versions:

|  |
| --- |
| Cellranger from 10x Genomics version 6.0.0 (RRID:SCR_017344). |
| subset-bam from 10x Genomics version 1.1.0 (RRID:SCR_023216). |
| bamtofastq from 10x Genomics version 1.4.1 (RRID:SCR_023215). |
| STAR aligner version 2.7.8a (RRID:SCR_004463). |
| SAMtools version 1.18 (RRID:SCR_002105). |
| TEcount version 2.2.3 (RRID:SCR_023208). |
| featureCounts subread version 1.6.3 (RRID:SCR_012919). |
| Seurat version 5.0.2 (RRID:SCR_007322). |

TrusTEr was ran using unique mapping (unique = True) and accounting for genes (include_genes = True). SizeFactors for each pseudocluster were calculated using gene counts (DESeq2) and used to normalize pseudobulked TE expression.

## Data availability

All data needed to evaluate the conclusions in the paper are present in the paper and/or the Supplementary Materials. The sequencing data presented in this study has been deposited at the GEO superserie GSE283600. Proteomic data has been deposited to the ProteomeXchange Consortium via the PRIDE partner repository^84^ with the identifier PXD059661. Original code has been deposited at GitHub and is publicly available at https://molecular-neurogenetics.github.io/truster/ (DOI:10.5281/zenodo.14362613) and git@github.com:Molecular-Neurogenetics/L1CRISPR_Adami_2025.git (DOI:10.5281/zenodo.14656834).

The data, code, protocols, and key lab materials used and generated in this study are listed in a Key Resource Table which will be available upon publication.

## IP rights notice

For the purpose of open access, the authors have applied a CC_BY public copyright license to the Author Accepted Manuscript (AAM) version arising from this publication.

## Acknowledgements

We would like to thank D. Trono and R. Foray for excellent comments on the manuscript and S. Henikoff for providing reagents. We also thank U. Jarl, A. Hammarberg for technical assistance. We acknowledge and thank J. Hansson and S. Ghosh for producing the mass spectrometry data and contributing with their expertise in proteomics data. We acknowledge Clinical Genomics Lund, SciLifeLab, and Center for Translational Genomics (CTG) Lund University for providing expertise and service with sequencing and analysis. We are grateful to all members of the Jakobsson laboratory.

## Author contributions

Design and interpretation: All authors. Conceptualization: A.A., R.G., and J.J. Experimental research: A.A., P.G., P.A.J, F.D., S.K., L.C-V, D.A.M.A., J.G.J,. Bioinformatics: R.G., P.G., Y.S., O.T., and C.H.D. Material, reagents, and expertise: A.K., M.H., and R.A.B.. Writing—original draft: A.A., R.G., and J.J. Writing—review and editing: All authors.

## Supplementary figures

**Supplementary Figure 1.**
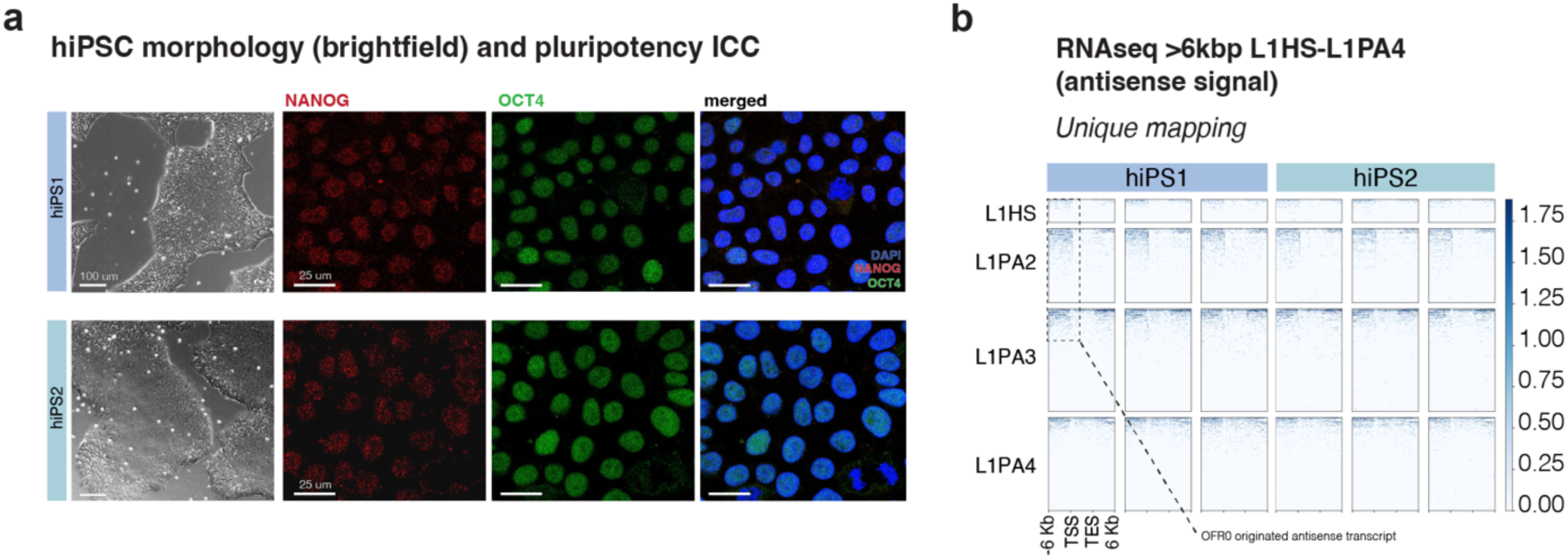
L1s are highly expressed in hiPSCs. a) Antisense expression of FL-L1s in two hiPSC lines. n= 3 technical replicates (normalized by RPKM). b) Immunocytochemistry of the pluripotency marker NANOG (red), OCT4 (green) in hiPSCs. DAPI nuclear staining (blue) is included in the overlay.

**Supplementary Figure 2.**
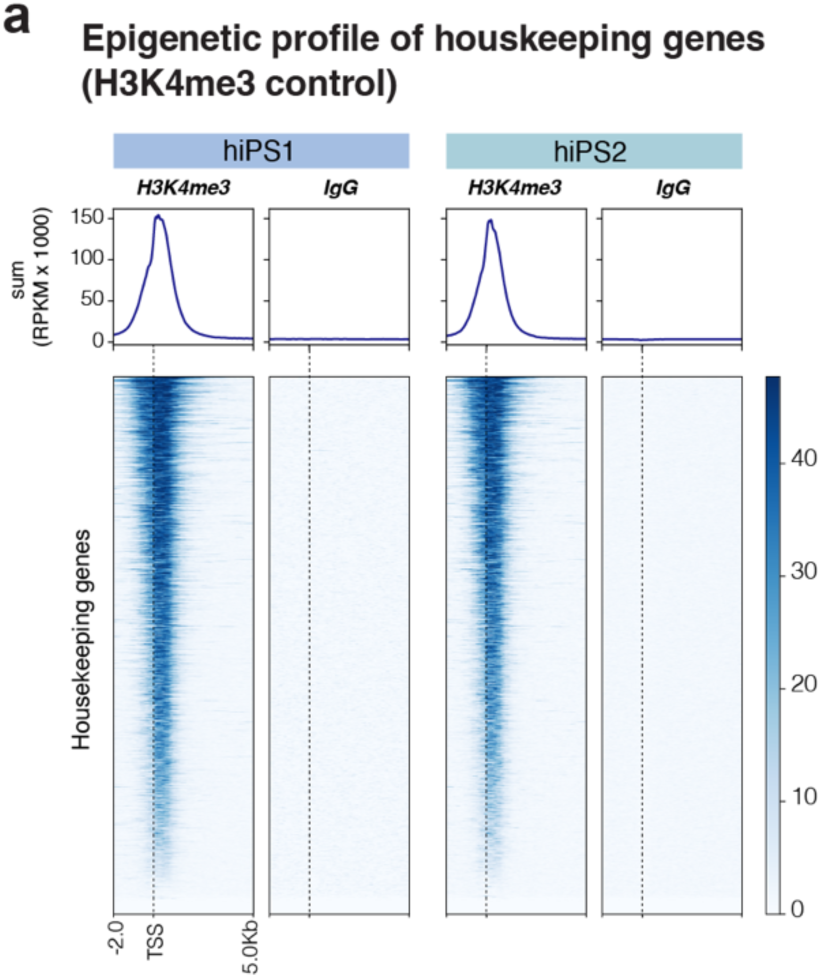
H3K4me3 CUT&RUN control. a) CUT&RUN profiles of the active histone mark H3K4me3 over housekeeping genes (RPKM normalised). TSS = transcription start site. Profile plot at the top showing the summed signal.

**Supplementary Figure 3.**
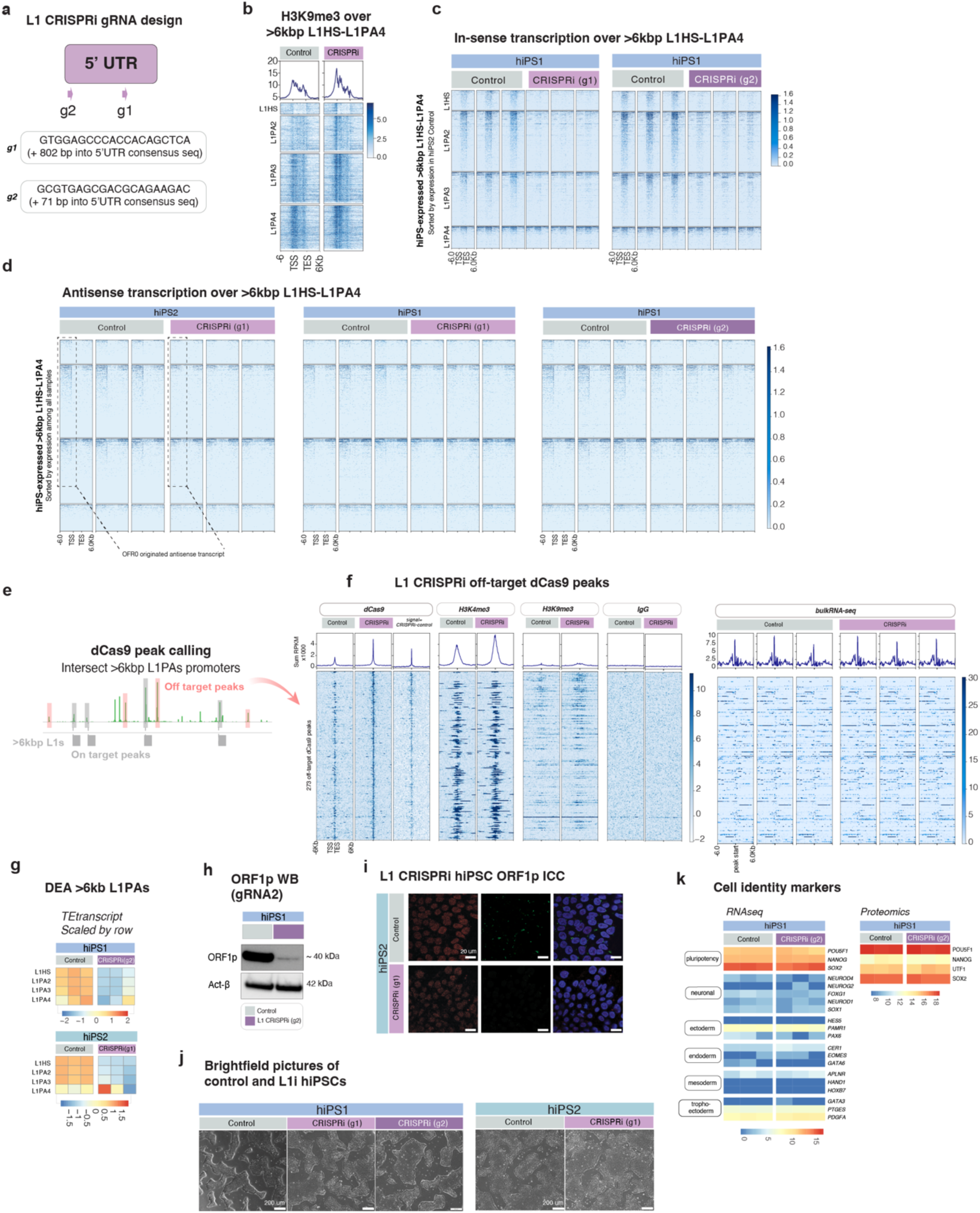
CRISPRi-based silencing of L1s in hiPSCs does not affect human pluripotency. a) Schematic of the gRNA target sites within the FL-L1. gRNAs designed to target the 5’ UTR of L1 consensus sequence. b) CUT&RUN analysis of H3K9me3 over young, FL-L1s in control vs. L1 CRISPRi hiPSCs (RPKM normalized). Profile plot on top showing summed signal. c) Bulk RNA seq data showing the normalized expression of uniquely mapped, FL-L1s in L1-CRISPRi vs. control hiPSCs (all tracks RPKM normalized). d) Antisense expression (RPKM) of FL-L1s in control vs. L1 CRISPRi hiPSCs. Two hiPSC cell lines, two guide RNAs, three technical replicates each. d) Bioinformatical rationale to identify off-target effects based on dCas9 CUT&RUN. e) CUT&RUN showing dCas9 (gRNA1 - Control), H3K4me3, H3K9me3, IgG, and RNA-seq in control vs. L1 CRISPRi hiPSCs over off-target peaks (dCas9 peaks not overlapping with FL-L1s promoters) (all tracks RPKM normalized). g) Expression of L1 families analysed using TEtranscripts in L1-CRISPRi vs. control hiPSCs in second gRNA and cell line (heatmap showing normalized expression, scaled by row) h) Western blot (WB) of ORF1p (top) and actin-β (bottom) in L1-CRISPRi using gRNA2 vs. control hiPSCs. j) Immunostaining on hiPSC2 of pluripotency marker NANOG (red) and L1-derived protein ORF1p (green) in L1-CRISPRi vs. control hiPSCs. DAPI nuclear staining in blue. i) Mass spectrometry (MS) data showing changes in ORF1p levels in L1-CRISPRi hiPSCs. j) Brightfield imaging of L1-CRISPRi vs. control hiPSCs in both cell types and gRNAs. k) Left panel: Heatmap showing log2 normalized expression (RNA-seq) of pluripotency and differentiation markers in L1-CRISPRi using gRNA2 vs. control hiPSCs. Right panel: Heatmap of MS data showing expression of pluripotency markers in L1-CRISPRi using gRNA2 vs. control hiPSCs (heatmap showing log2 normalized expression).

**Supplementary Figure 4.**
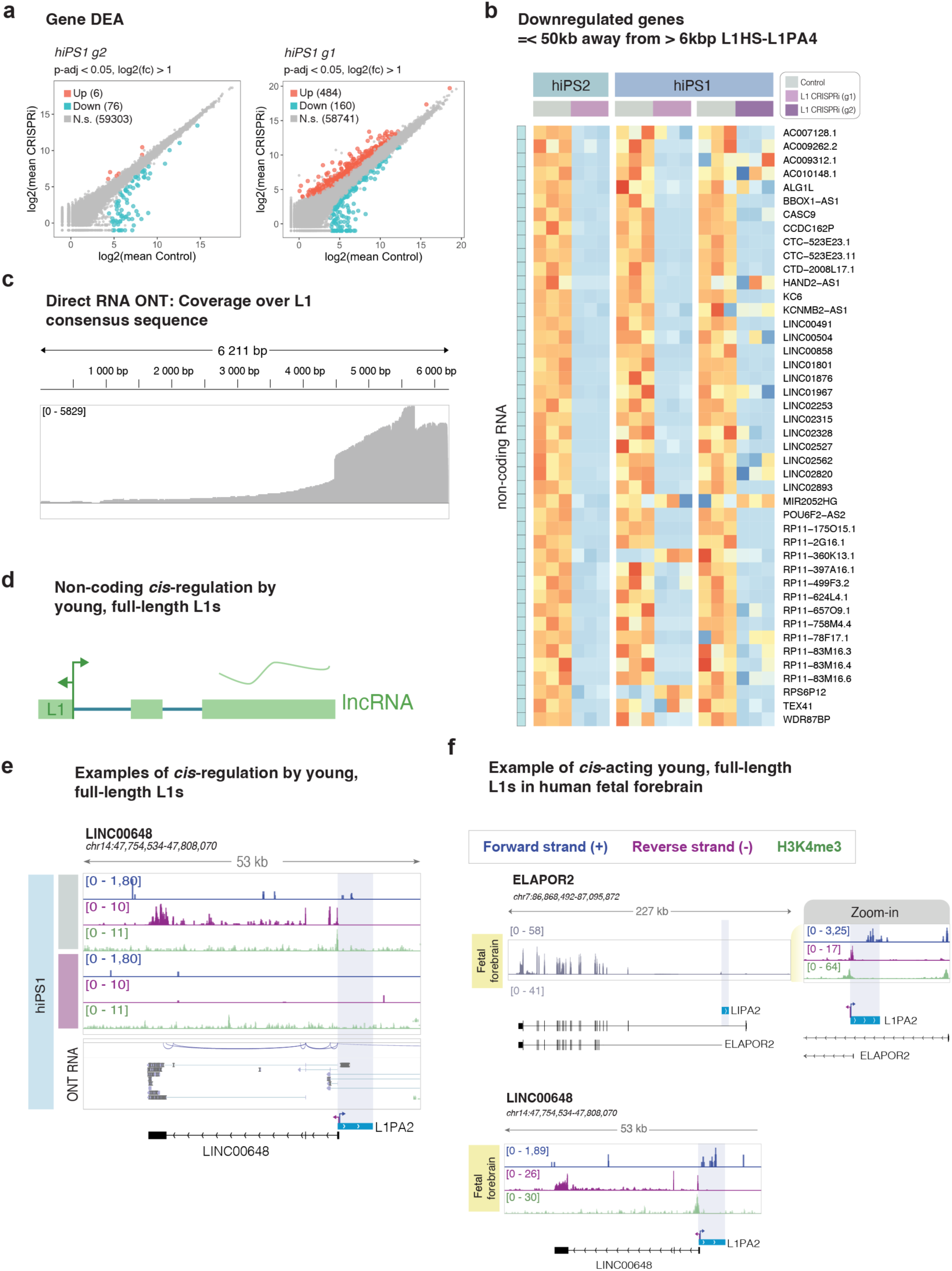
L1s influence the expression of protein coding genes and long non-coding RNAs in *cis*. a) Scatterplot showing mean gene expression in L1-CRISPRi (y-axis) and control (x-axis) hiPSCs using second gRNA and cell line. Plot showing summarized results of differential expression analysis (DEA) (Bulk RNA-seq, n = 3 replicates / condition). Blue dots = significantly downregulated genes; red dots = significantly upregulated genes; grey dots = non-significant (DESeq2: Wald test; p-adj <0.05; log2(foldChange) >1). b) Heatmap showing all the normalized expression of downregulated non-coding genes that overlap with a FL-L1 (n= 3 replicates / condition, heatmap showing log2 normalized expression). c) Direct RNA ONT reads mapped to L1 consensus sequence. d) Schematic showing the expression of a L1-driven long non-coding RNA. e) Genome browser tracks showing the expression of an L1-driven long non-coding RNA in hiPSCs, and its silencing upon L1-CRISPRi. From top to bottom: control hiPSC bulk RNA-seq data (dark blue: forward transcription, purple: reverse transcription), H3K4me3 CUT&RUN (green, control vs L1-CRISPRi hiPSCs), L1-CRISPRi hiPSC bulk RNA-seq data, H3K4me3 CUT&RUN, and control hiPSC ONT direct RNA reads from control hiPSCs. f) Top: genome browser tracks showing the expression of ELAPOR2 in human fetal forebrain. From top to bottom: bulk RNA-seq data in grey (left), zoom-in (right) showing bulk RNA-seq data (dark blue: forward transcription, purple: reverse transcription), and H3K4me3 CUT&RUN (green). Bottom: genome browser tracks showing the expression of LINC00648 in human fetal forebrain. From top to bottom: bulk RNA-seq data (dark blue: forward transcription, purple: reverse transcription), and H3K4me3 CUT&RUN (green) (all tracks normalized by RPKM). Data from Garza *et al.* (2023)26.

**Supplementary Figure 5.**
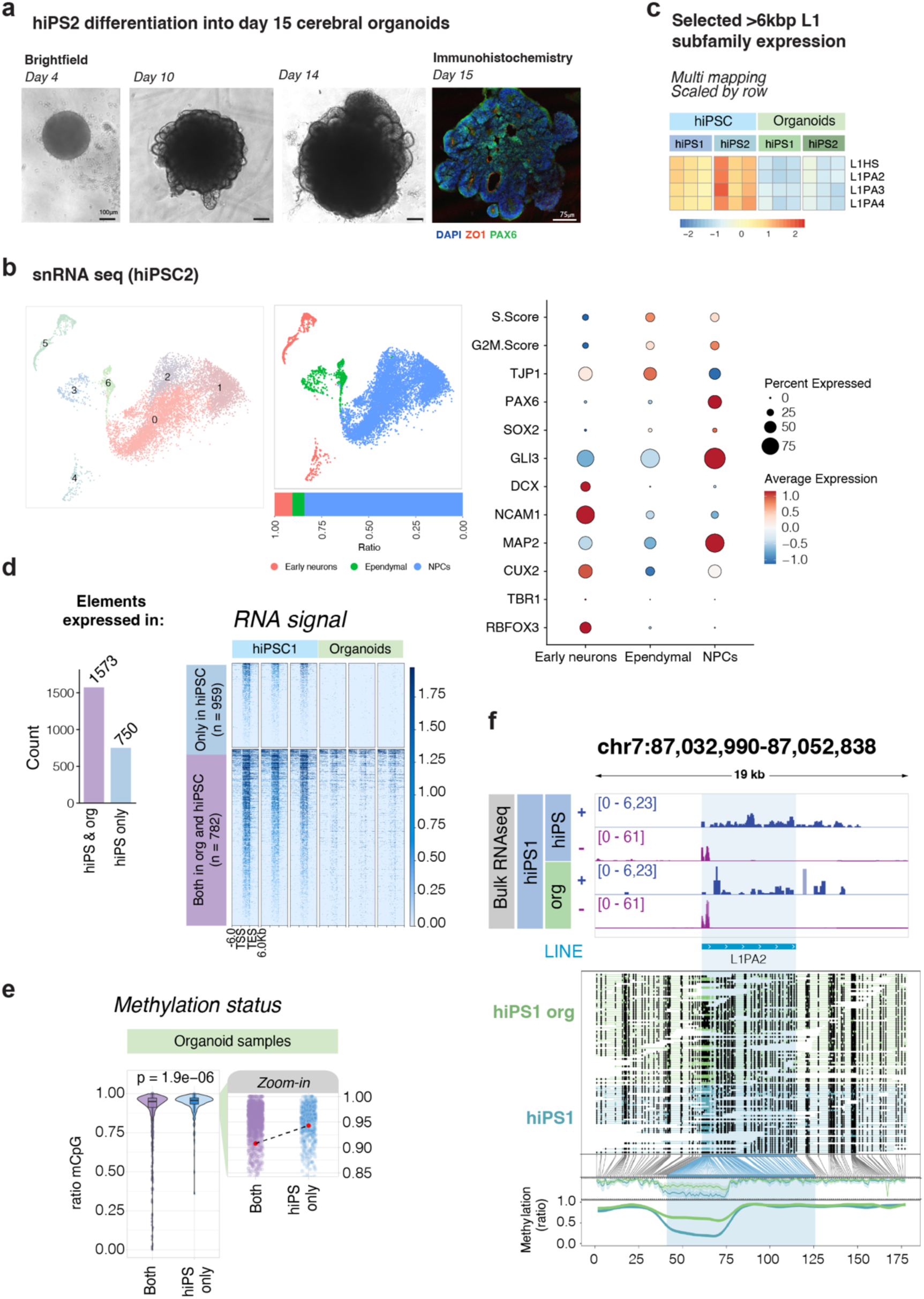
Expression of evolutionary young full-length L1s in cerebral organoids. a) Brightfield images of differentiating cerebral organoids (hiPSC2) at different time points and immunohistochemistry on day 15 (hiPSC2) organoids for ZO1 (red) and PAX6 (green). DAPI in blue. b) Left: UMAP showing clusters found in day 15 hiPSC2 unguided cerebral organoids. Middle: UMAP showing cell types found in day 15 hiPSC2 unguided cerebral organoids. Bar plot showing the percentage of the cell type composition of the cerebral organoids. Right: dot plot displaying selected neuronal and NPC markers used to characterize the cell clusters (dot size showing the percentage of cells expressing the gene, color indicates average expression in each cell type). c) Expression of L1 families analyzed using TEtranscripts in L1-CRISPRi vs. control organoids and hiPSCs (heatmap showing normalized expression, scaled by row). d) Left: Barplot showing the number of expressed FL-L1 in hiPSC and day 15 cerebral organoids (purple), and hiPSC only (blue). Right: Bulk RNA seq data showing the normalized expression (RPKM) of FL-L1 in hiPSC and day 15 cerebral organoids (purple, bottom), and hiPSC only (blue, top) in hiPSC1 and day 15 hiPSC1-derived cerebral organoids (n = 3 replicates). e) Violin plots of the methylation status over the promoter of FL-L1 in hiPSC and cerebral organoids (purple), and hiPSC only (blue). Zoom in panels indicating mean methylation levels (red dot). f) Genome browser tracks showing normalized (RPKM) of L1s in hiPSCs and day 15 cerebral organoids (dark blue: forward transcription, purple; reverse transcription) and ONT DNA reads across day 15 organoids and hiPSCs, black dots indicating methylated CpGs, and methylation coverage of the L1 elements at the bottom (green: organoids, blue: hiPSCs).

**Supplementary Figure 6.**
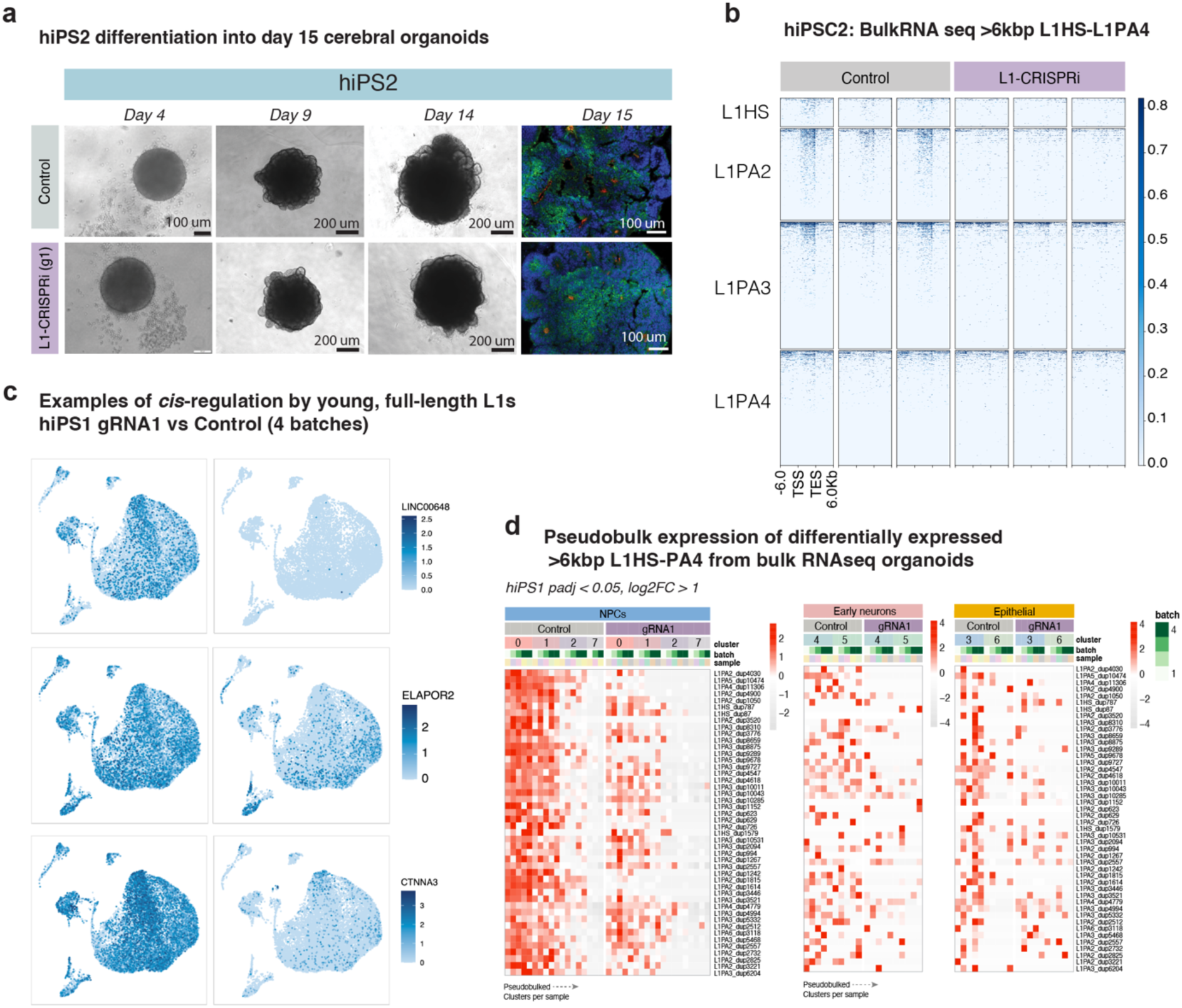
a) Brightfield images of differentiating cerebral organoids (L1-CRISPRi and control hiPSC2) at different time points and immunohistochemistry on day 15 L1-CRISPRi and control hiPSC2 organoids for ZO1 (red) and PAX6 (green). DAPI in blue. b) Normalised expression (RPKM) of uniquely mapped evolutionary young (L1HS-L1PA4), >6kbp L1s in control vs. L1-CRISPRi unguided cerebral organoids at day 15 of differentiation using hiPSC2. c) Heatmap showing differentially expressed FL-L1s from bulk RNA-seq of L1-CRISPRi (gRNA1 vs control hiPSC1) (DESeq2: Wald test; padj < 0.05, log2FC < -1) in snRNA-seq pseudobulks (n = 8 replicates (4 = gRNA1, 4 = control), in four organoid batches). Expression normalized by gene sizeFactors as calculated by DESeq2 (i.e. median of ratios) using pseudo bulked gene expression. d) UMAPs showing L1-derived gene expression

